# Impact of wildfire smoke and diesel exhaust on inflammatory response in aging human microglia

**DOI:** 10.1101/2024.12.20.629570

**Authors:** Carla Cuní-López, Mei Fong Ng, Romal Stewart, Laura A. Milton, Fazeleh Etebar, Yifan Sun, Emily Vivian, Tam Hong Nguyen, Patrick F. Asare, Michelle K. Lupton, Tara L. Roberts, Zoran Ristovski, Sandra Hodge, Paul N. Reynolds, Anthony R. White, Hazel Quek

## Abstract

**Background:** Air pollution, particularly from Diesel Exhaust Particles (DEP) and Wildfire Smoke (WFS), is increasingly recognised as a significant driver of neuroinflammation linked to brain diseases. However, the role of microglia in mediating these neuroinflammatory responses remains poorly understood. This study aimed to investigate the effects of air pollution on monocyte-derived microglia-like cells (MDMi) from both young (< 40 years of age), and (older > 60 years of age) healthy individuals, focusing on immune response, cytokine secretion, nitrosative stress, and phagocytic activity.

**Results:** Our study demonstrated that DEP and WFS extract (WFSE) significantly upregulated expression of the oxidative stress marker, heme-oxygenase-1 (HO-1) in MDMi after 24 hr, with levels normalising by 96 hr, indicating a transient oxidative stress response. Both DEP and WFSE elicited distinct inflammatory cytokine profiles. DEP induced a rapid response, increasing TNF-α, IL-6, IL-23, and IL-33 within 2 hr in young MDMi and 24 hr in aged MDMi. In contrast, WFSE triggered a delayed but sustained inflammatory response, with TNF-α, IFN-γ, IL-23, and IL-33 levels persisting at 96 hr in aged MDMi, highlighting an age-related vulnerability to air pollutant-induced inflammation.

Both pollutants activated p38, ERK, and NF-κB pathways, with p38 activity resolving by 96 hours and ERK activation persisting, reflecting their distinct roles in cellular stress and inflammation. NF-κB p65 nuclear translocation, observed at 24 hours, highlighted its critical role in cytokine release and inflammation following exposure to DEP and WFSE. This is the first report of NF-κB activation in human microglia exposed to air pollutants.

**Conclusions:** These results highlight the distinct and potentially harmful effects of DEP and WFSE on immune and inflammatory responses in MDMi, particularly in ageing populations, with significant implications for brain health. DEP triggers acute oxidative stress and inflammatory responses, while WFSE induces more prolonged effects, especially in aged microglia. Both pollutants activate the MAPK and NF-κB pathways and exhibit unique cytokine profiles, underscoring their overlapping yet distinct mechanisms of action. These findings advance our understanding of air pollutant-induced neuroinflammation and its contribution to neurodegeneration, providing a foundation for developing targeted interventions to mitigate the neurotoxic effects of air pollution.

## Background

Air pollution is a pressing global public health concern, especially in the midst of global warming and the rapidly ageing global population [1, 2]. Major cities worldwide are affected by significant levels of traffic-related air pollution. Despite efforts to mitigate emissions, urban and some regional areas remain plagued by high concentrations of pollutants such as nitrogen dioxide, particulate matter (PM), and carbon monoxide, largely stemming from vehicular exhaust. In particular, diesel engine pollution is a major factor due to higher levels of PM, and gaseous emissions than other fuel types [3–5]. These pollutants pose severe health risks, and premature deaths. Various measures are being pursued to address this pressing issue, however, some of the improvements are now being lost to elevated levels of wildfire smoke pollution [6]. Wildfire smoke also contains a complex mixture of harmful pollutants, including PM, carbon monoxide, volatile organic compounds, and hazardous chemicals. These pollutants can travel long distances from origin, impacting air quality and posing health risks to populations over large areas [7]. Climate change is leading to increased frequency and severity of wildfires, and associated smoke. In-depth studies have established a correlation between air pollutants from both vehicular and wildfire sources and detrimental health consequences, predominantly manifesting as respiratory and cardiovascular disorders [8]. Interestingly, increasing incidence of neurodegenerative diseases, including Alzheimer’s disease [9–12] and Parkinson’s disease [13] is observed in areas with high levels of air pollution, and this is supported by epidemiological studies showing links between air pollution and incidence of neurodegenerative diseases.

In line with this, neuropathological hallmarks and biomarkers associated with neuroinflammation have been found in post-mortem brains and cerebrospinal fluid of individuals exposed to high levels of air pollution [14–16]. Although the mechanisms of air pollution on brain function and disease are still not fully understood, increasing evidence supports the toxic effects of PM in various CNS diseases.

PM, a significant component of air pollutants, is generally categorised into three size fractions; coarse (PM 10: 2.5-10 μm), fine (PM 2.5: < 2.5 μm) and ultrafine (PM 0.1 < 0.18 μm) [17, 18]. PM can penetrate the respiratory tract and potentially cross the blood-brain barrier, reaching the central nervous system (CNS) and triggering neuroinflammation and neurodegeneration [19, 20]. Notably, studies have shown that fine or ultrafine PM particles can elicit differential immune responses in the brain, suggesting that PM size may modulate the resulting immune response [21].

Interestingly, PM exposure can activate microglia, the immune cells of the CNS, indirectly through the peripheral immune system [22, 23] or directly through inhaled air [24], leading to the release of pro-inflammatory cytokines, and chemokines, as well as inducing DNA damage and oxidative stress [25–27]. Increased neuroinflammation has been observed to exacerbate neuronal and oligodendrocyte cell death [28–31], and increase protein misfolding and amyloid-beta plaques, which are hallmarks of Alzheimer’s disease [32]. Hence, elucidating the processes through which PM exposure contributes to neuroinflammation and neurodegeneration is essential in tackling the ramifications of air pollution on the neurological well-being of the growing elderly population.

Various non-particulate components of air pollution have also been implicated in neuroinflammation, contributing to a range of neurological disorders. Nitrogen dioxide (NO_2_), a common pollutant emitted by vehicles, has been linked to neuroinflammation and neurodegenerative diseases such as Alzheimer’s and Parkinson’s diseases [33, 34]. NO_2_ exposure has been associated with increased levels of pro-inflammatory cytokines and oxidative stress markers in the brain. Similarly, exposure to ozone (O_3_), a major component of wildfire smoke, has been shown to induce neuroinflammation and impair cognitive function [35, 36]. Studies have demonstrated that O_3_ exposure leads to elevated levels of inflammatory mediators in the brain and disrupts the blood-brain barrier, facilitating the infiltration of immune cells into the central nervous system [36]. Additionally, volatile organic compounds (VOCs) present in air pollution, such as benzene and toluene, have been implicated in neuroinflammatory responses and neurotoxicity [37, 38]. These findings highlight the detrimental impact of various non-particulate components of air pollution on neuroinflammation and highlight the urgent need for further studies on the impact of air pollution on neuroinflammatory contributions to neurodegenerative disorders.

Previous studies exploring the impact of air pollutants on microglia in the context of neuroinflammation have primarily concentrated on rodent models while providing only a restricted analysis of immortalised human microglia [20]. Therefore, this study aimed to fill this gap by utilising human monocyte-derived microglia-like cells (MDMi) to evaluate the effects of air pollution from wildfire smoke and diesel engine emission. With its physiological relevance, the MDMi model can effectively represent individual demographics (including age and sex), clinical variables (such as disease severity), and risk factors (encompassing lifestyle and environmental aspects), making it an appropriate tool for investigating the fundamental mechanisms through which air pollution exposure influences microglial cells in a research setting [39–41]. Understanding these processes is essential in addressing the neurological impacts of air pollution on the ageing population.

## Methods

### Recruitment of young and aged blood donors

Peripheral blood samples were collected from two cohorts of healthy individuals with no reported clinical symptoms: young donors (n = 6, aged < 40 years) and aged healthy controls (HC; n = 6, aged > 60 years). The aged cohort was recruited through the Prospective Imaging Study of Aging: Genes, Brain, and Behaviour (PISA)[42]. Both cohorts were recruited at QIMR Berghofer Medical Research Institute, Queensland, Australia. A summary of the demographics of the donor cohorts is shown in Supplementary Table 1. All research adhered to the ethical guidelines on human research outlined by the National Health and Medical Research Council of Australia (NHMRC). Ethical approval was obtained from QIMR Berghofer Medical Research Institute (P2197). All participants provided informed consent before participating in the study.

### PBMC isolation and MDMi differentiation

Peripheral blood mononuclear cell (PBMC) isolation and MDMi differentiation were performed according to established protocol [43]. Briefly, PBMCs were plated onto a Matrigel coated plate and cultured with RPMI-1640 Glutamax media supplemented with 1% penicillin/streptomycin (P/S), 0.1 µg/mL interleukin (IL)-34 and 0.01 µg/mL granulocyte-macrophage colony-stimulating factor (GM-CSF) (ThermoFisher) for 14 days. On day 14, MDMi were treated with air pollutants and harvested at designated time points. MDMi cultures have been extensively characterised and previously described by us and others [41, 44–47].

### Generation of wildfire smoke extract (WFSE)

A 100% WFSE stock was kindly generated and provided by the Chronic Lung Disease Research Laboratory at The University of Adelaide as previously described [48, 49]. Briefly, 2 g of relevant Australian wildfire susceptible plant material (*Acacia melanoxylon, Acacia vestita, Eucalyptus camaldulensis* and *Eucalyptus globulus* in equal masses) was burnt and bubbled through 20 mL of Opti-MEM media, before the pH was adjusted to neutrality and stored at −80 °C.

### Generation of diesel exhaust particles (DEP)

Nanometre-sized DEP were used as a model for ultrafine PM and was kindly provided by the International Laboratory for Air Quality and Health at the Queensland University of Technology. The exhaust samples were collected using a 4-cylinder Perkins diesel engine coupled with an eddy current dynamometer to maintain steady-state conditions. The engine was warmed up at 50% load and 1,500 rpm for 20 minutes, and then operated at 1,500 rpm and 75% load during the experiments. Exhaust samples were analysed using an ejector diluter (Dekati), gas analyser, and DMS500 particle size analyser [50]. Carbon dioxide (CO_2_) measurements were taken using a CAI600 series NDIR analyser for raw exhaust, and a Sable CO_2_ meter after the diluter to calculate the dilution ratio. The exhaust samples were collected in serum-free RPMI-1640 GlutaMax medium supplemented with 2% P/S using a fritted glass impinger at a flow rate of 1 L/min [51]. The sampling time for the media samples was 45-50 min. The collected particle mass was determined from DMS500 data using a re-inversion tool in DMS data analysis software and corrected for the known sampling efficiency of the impingers[51]. This concentration was selected based on *in vivo* models as an appropriate *in vitro* surrogate for DEP concentrations (5-50 μg/L) that reach the brain [52].

### Lactate dehydrogenase (LDH assay)

Cell media of MDMi treated with WFSE and DEP and relevant assay controls (media alone and positive lysed control) were harvest and centrifuged at 500 *g* for 5 min and stored at −80°C until use. LDH was measured using the CytoTox-ONE membrane integrity assay (Promega) according to manufacturer’s instructions. OD 560 nm was measured using a BioTek Synergy Neo2 spectrophotometer. LDH release was calculated as follows:

(Experimental - Culture medium background) / (Maximum LDH release - Culture medium background).

### Endotoxin testing of WFSE and DEP

Cell media treated with WFSE or DEP was harvested and tested for endotoxin using a chromogenic endotoxin kit (ThermoFisher) as per the manufacturer’s instructions. OD 405 nm was measured using a BioTek Synergy Neo2 spectrophotometer. A standard curve was used to determine the endotoxin concentration, which was less than the limit of detection (0.01 EU/mL).

### Cytokine and chemokine profiling

Cell media of MDMi treated with WFSE or DEP was harvested at respective time points, centrifuged at 400 *g* for 5 min to remove cell debris and stored at −80°C until use. Cytokine and chemokine concentrations were measured using a bead-based multiplex LEGENDplex™ Human Inflammation Panel 1 (BioLegend) kit as per the manufacturer’s instructions. The panel includes 13 cytokine/chemokines: IL-1β, IFN-α2, IFN-γ, TNF-α, MCP-1 (CCL-2), IL-6, IL-8 (CXCL8), IL-10, IL-12p70, IL-17A, IL-18, IL-23, and IL-33. Sample acquisition was performed in duplicates using a flow cytometer, and the data were analysed using Qognit (BioLegend, USA) specified at pg/ml values. Cytokines below the limit of detection were excluded from the analysis indicated in the respective figure legends.

### Measurement of nitrate concentration

Nitrite, a by-product of NO oxidation, was measured using the Griess method. Cell media of MDMi treated with WFSE or DEP was harvested after 24 hr and levels of nitrite were determined using the Griess reagent (Promega) according to the manufacturer’s instructions. To measure nitrite, 1:1 ratio of Griess reagent (sulphanilamide solution) and cell media were incubated at room temperature for 10 min. After incubation, equal volumes of N-1-napthylethylenediamine dihydrochloride (NED) was added and incubated for another 10 min. The nitrite concentration was determined from a sodium nitrite standard curve, with a lower limit of detected of 2.5 µM. OD 540 nm was measured using a BioTek Synergy Neo2 spectrophotometer.

### Phagocytosis assay

MDMi at Day 14 were treated with 1% WFSE or 5.7 mg/L DEP in RPMI + GlutaMax media supplemented with 1% P/S for 1 hr, along with an untreated control. Red fluorescent pHrodo dye labelled-*Escherichia coli* (*E. coli*) BioParticles (ThermoFisher) were added to a final concentration of 20 µg/mL. The plate was imaged using the IncuCyte SX5 system (Essen Bioscience), where sixteen phase-contrast and red fluorescence images were taken per well, every hour for 12 hr. As the pHrodo dye fluoresces red inside the acidic phagosome, the IncuCyte software was utilised to quantify the total red area in each image to measure uptake. Red area was divided by the cell area for normalisation. Area under the curve (AUC) was calculated for each technical replicate for statistical analysis.

### Real-time PCR

Cells were incubated with 200 µL of TRIzol (ThermoFisher) for 5 min before being collected and stored at −80°C. RNA was extracted via the TRIzol-chloroform method. RNA concentration and quality was quantified using the NanoDrop2000. cDNA synthesis was performed using a SensiFast cDNA synthesis kit (Bioline). Samples were diluted and mixed with SensiFAST Sybr Lo-Rox master mix before loading as triplicates for qRT-PCR. qRT-PCR was performed using the Applied Biosystems ViiA 7 system (ThermoFisher). Delta cycle threshold (ΔCT) values were normalised to *18S* and used to determine relative gene expression between samples. Samples with CT values more than two times the standard deviation of the average of each reference gene were excluded from the analysis. The primers used in this study are found in the Supplementary Table 2. Melting curve analyses confirmed a single melting curve peak for all primers.

### Immunofluorescence

Cells were plated onto 8-well chamber slides (Ibidi), treated with WFSE or DEP at multiple time points and immunofluorescence staining was performed. Briefly, cells were fixed for 15 min with 4% paraformaldehyde in PBS. Permeabilisation of PFA fixed samples was performed with PBS containing 0.3% Triton-X 100 (Sigma-Aldrich). Samples were blocked with PBS containing 2.5% bovine serum albumin (BSA) (Sigma-Aldrich) in PBS. NF-κB p65 antibody (sc-372, 1:300) was incubated overnight at 4°C. Cells were washed thrice with 0.1% Triton-X in PBS followed by incubation with secondary antibodies and 1µg/mL Hoechst (nuclear dye) (Sigma-Aldrich) for 2 hr at room temperature. Antibody specificity was confirmed by performing secondary antibody only controls. Images were captured with a confocal laser scanning microscope (20x NA0.8, LSM-780, Carl Zeiss) with all settings kept consistent during acquisition.

### NF-κB p65 nuclear translocation

To assess NF-κB p65 nuclear translocation, we measured mean nuclear and cytoplasmic fluorescence intensities in NF-κB p65-stained cells. Nuclear masks were generated by manually delineating nuclei based on Hoechst staining, which were then added to the region of interest (ROI) manager in ImageJ. The nuclear mask was subtracted from the NF-κB p65 channel to create a cytoplasmic ROI mask, ensuring only non-nuclear areas contributed to the cytoplasmic intensity measurement. Mean fluorescence intensities of nuclear and cytoplasmic ROIs were recorded for each cell, and nuclear localisation was calculated as follows:

% Nuclear = (Total Nuclear Intensity/ Total Cytoplasmic Intensity + Total Nuclear Intensity) * 100

### Western blot

MDMi cells were treated with 1% WFSE or 5.7 mg/L DEP for 24 hr and 96 hr. On the day of harvest, the cells were rinsed with ice-cold PBS and lysed with radioimmunoprecipitation buffer assay (RIPA) containing protease and phosphatase inhibitor cocktail (Roche; #05892791001) and incubation on ice for 10 min. After, lysed cells were centrifuged at 14,000 rpm, 4°C for 30 mins. Supernatants were collected and stored at −80°C until ready for protein determination using the Pierce™ BCA Protein Assay Kit (Thermo Fisher Scientific; #23225) according to manufacturer’s instructions.

Next, 30 μg of total cellular lysates were resolved in Bolt 4-12% Bis-Tris plus mini protein gels (Thermo Fisher Scientific) by SDS-PAGE followed by transferring onto methanol-activated Immobilon-PSQ PVDF transfer membrane (Merck). Membrane was blocked with 5% skimmed milk in Tris-buffered saline plus 0.1% Tween-20 (TBS-T) for 1 hr at RT and further incubated with the following antibodies: anti-HO-1 (#5853), anti-phospho-ERK1/2 (#9101), anti-ERK1/2 (#9102), anti-phospho-p38 (#9211), anti-p38 (#9212) and anti-β-actin (loading control; #4970, 1:5000) (Cell Signalling Technology) at 4°C overnight, in 5% BSA in TBS-T. All primary antibodies were used at 1:1000 unless otherwise specified. The following day, the membrane was washed with TBS-T and subsequently probed with respective secondary antibodies conjugated with horseradish peroxidase (1: 5000) in 5% skimmed milk in TBS-T for 1 hr at RT. After washing with TBS-T, the proteins were detected using SuperSignal^TM^ West Pico Plus Chemiluminescent substrate (Thermo Fisher Scientific) with iBright™ CL1500 Imaging System (Invitrogen). Assays were repeated twice, each time with an independent individual (n = 2). Quantification of protein bands were performed using FIJI ImageJ Version 1.49e. The relative intensity of target proteins was calibrated against β-actin while phospho-specific proteins were further normalised to total of respective protein to calculate fold change.

### Statistical analysis

All data was analysed with GraphPad Prism version 9 and is represented as mean ± standard error of the mean (SEM). Comparisons between two groups were completed with an unpaired student’s *t*-test if data is normal or Mann-Whitney *U* when normality assumptions were not met. Comparisons between three or more groups were analysed with a one- or two-way ANOVA followed by post-hoc tests. Data are presented as mean ± SEM or mean ± SD and and p ≤ 0.05 was considered significant. Statistical significance was determined as *p < 0.05, **p < 0.01, ***p < 0.001, ****p < 0.0001, as detailed in figure legends.

## Results

### Establishing non-toxic air pollutant concentrations to investigate immunomodulatory responses in MDMi cultures from young donors

MDMi were derived from blood monocytes through a 14-day trans-differentiation process using growth factors IL-34 and GM-CSF (**Fig. 1A**) [43]. Our previous work has established that MDMi cultures differ significantly from monocytes, exhibiting brain microglia-like features, including distinct single-cell transcriptomic signatures, morphology, and functionality (**Fig. 1B-C**) [53]. Building upon this foundation, the present study investigates the immunomodulatory impacts of air pollution specifically DEP and WFSE on the peripheral-brain axis using MDMi cultures (**Fig. 1D**).

**Figure 1.**
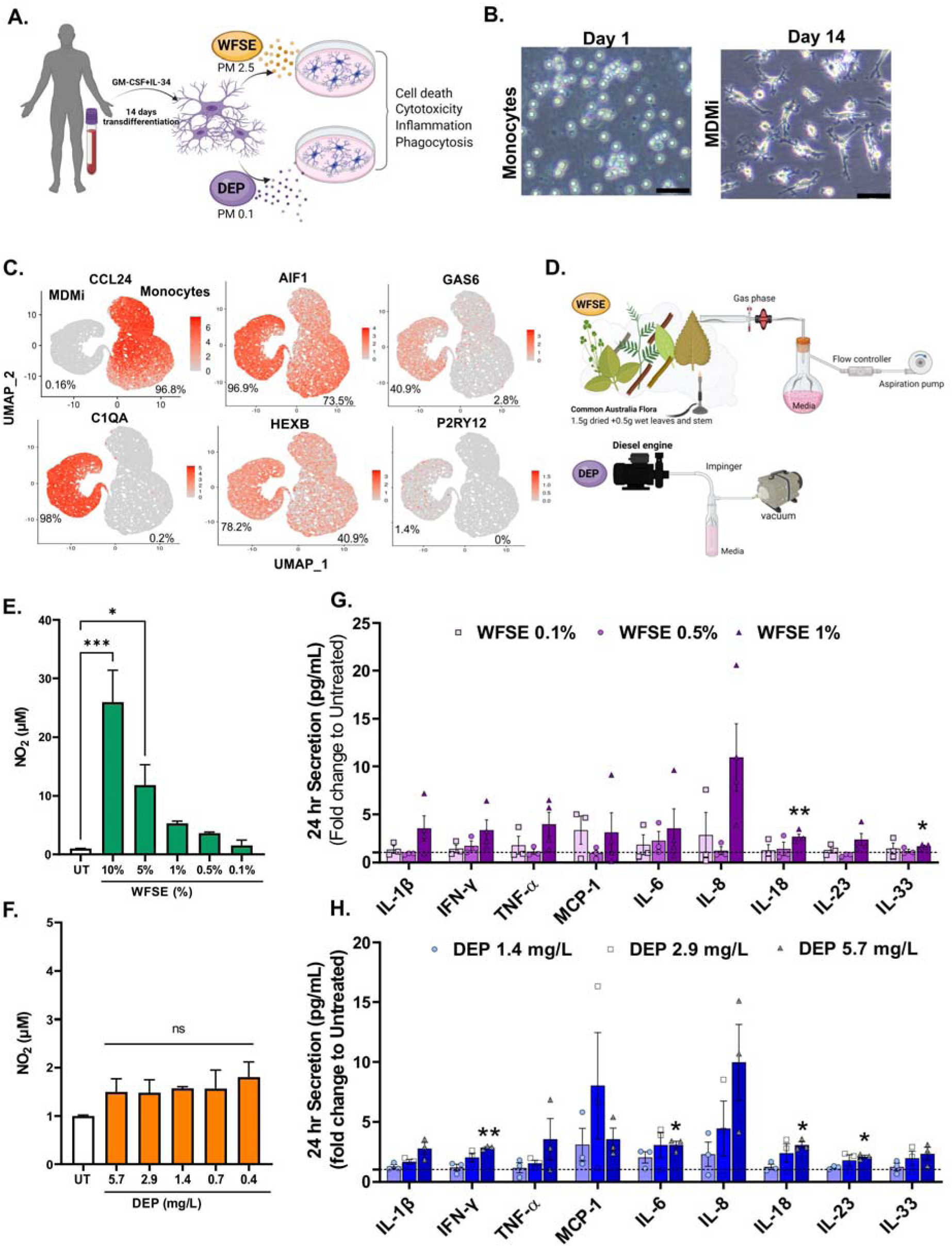
Assessment of cytotoxicity and immune response of WFSE and DEP on MDMi. **(A)** Blood-derived monocytes were differentiated into microglia-like cells (MDMi) over a 14-day period. Following differentiation, MDMi cells were exposed to WFSE or DEP for 24 hr, and inflammatory outcomes were assessed, including cell death, cytotoxicity, and inflammatory marker expression. **(B)** Phase-contrast images depict monocytes on day 1 of differentiation (left), showing transition to microglia-like morphology by day 14 (right). **(C)** The microglial identity of MDMi were evaluated by comparing their mRNA expression profiles to those of blood monocytes, using data derived from previously published single-cell sequencing study [53]. Each dot represents an individual cell within the model, with higher gene expression levels indicated by greater intensity of red coloration. **(D)** Schematic of WFS and DEP extracts used in the study. **(E, F)** Nitric oxide secretion increased in MDMi exposed to **(E)** 10% and 5% WFSE, with no significant changes observed in **(F)** DEP-exposed MDMi over 24 hr. **(G-H)** Cytokine analysis of **(G)** WFSE- and **(H)** DEP-exposed MDMi demonstrated a dose-dependent inflammatory response. Data presented as mean ± SEM, with data points representing biological replicates (n = 3 donors). Statistical analysis between two groups was performed using Student’s *t* test and between multiple groups using one-way ANOVA. Statistical significance indicated as follows: *p < 0.05, **p < 0.01, ***p < 0.001. Scale bars represent 50 µm.

The concentrations of WFSE were selected based on prior studies showing that up to 10% WFSE is well-tolerated in THP-1 and monocyte-derived macrophages [48]. To ensure relevance to MDMi, we first conducted dose-response experiments to evaluate cytotoxicity and inflammation. MDMi derived from a cohort of young individuals (<40 years, n = 3) were exposed to WFSE (0.1%, 0.5%, 1%, 5%, and 10%) or DEP (0.4 mg/L, 0.7 mg/L, 1.4 mg/L, 2.9 mg/L, and 5.7 mg/L) for 24 hr.

Exposure to WFSE at 5% and 10% concentrations resulted in significantly elevated nitric oxide (NO) secretion (**Fig. 1E**), which correlated with increased lactate dehydrogenase (LDH) levels, indicating cytotoxicity (**Fig. S1A**). Additionally, WFSE exposure induced cell death, as evidenced by a marked increase in caspase 3/7-positive cells, with 80% and 20% of MDMi cells showing positivity at 10% and 5% WFSE, respectively, after 24 hr (**Fig. S1C-E**).

In contrast, MDMi exposed to DEP at all tested concentrations (0.4–5.7 mg/L) demonstrated low nitrite secretion (**Fig. 1F**) and maintained cell viability (**Fig. S1B, D-F**). Consequently, WFSE concentrations of 5% and 10% were excluded from further analysis due to their cytotoxic effects.

To identify non-toxic concentrations, subsequent experiments focused on WFSE at 0.1%, 0.5%, and 1% and DEP at 1.4 mg/L, 2.9 mg/L, and 5.7 mg/L. After 24 hr of exposure, media was collected and analysed using a 13-plex human cytokine inflammation panel. Four cytokines (IL-10, IL-12p70, IL-17A, IFN-α2) were below detection limits and excluded from analysis. A dose-dependent increase in cytokine secretion was observed with both WFSE and DEP exposure.

Notably, 1% WFSE exposure resulted in a significant upregulation of IL-18 and IL-33 compared to untreated controls (**Fig. 1G**). Similarly, exposure to the highest DEP concentration (5.7 mg/L) significantly upregulated several cytokines, including IFN-γ, IL-6, IL-18, and IL-23, compared to untreated controls (**Fig. 1H**). Based on these findings, 1% WFSE and 5.7 mg/L DEP were selected for further investigation. These concentrations were chosen based on two criteria: 1) the absence of cytotoxic effects and 2) their ability to induce a measurable immune response.

Finally, endotoxin testing of the WFSE and DEP extracts confirmed the absence of LPS contamination (limit of detection <0.01 EU/mL), indicating that the observed pro-inflammatory responses were driven directly by air pollutants rather than secondary bacterial contamination (**Fig. S1G).**

### Exposure to air pollutants promotes nitrosative stress without altering phagocytic capability in MDMi

Activated microglia release pro-inflammatory mediators, such as NO, in response to air pollution exposure, a process implicated in neurodegeneration [29, 54]. However, the effects of air pollution on microglia-associated nitrosative stress, particularly in the context of ageing, remain poorly understood. To investigate this, along with effects on cytokine release and phagocytic activity, young and aged MDMi were exposed to 1% WFSE or 5.7 mg/L DEP for 24 hr (acute exposure) and 96 hr (chronic exposure) (Fig. 2A).

**Figure 2.**
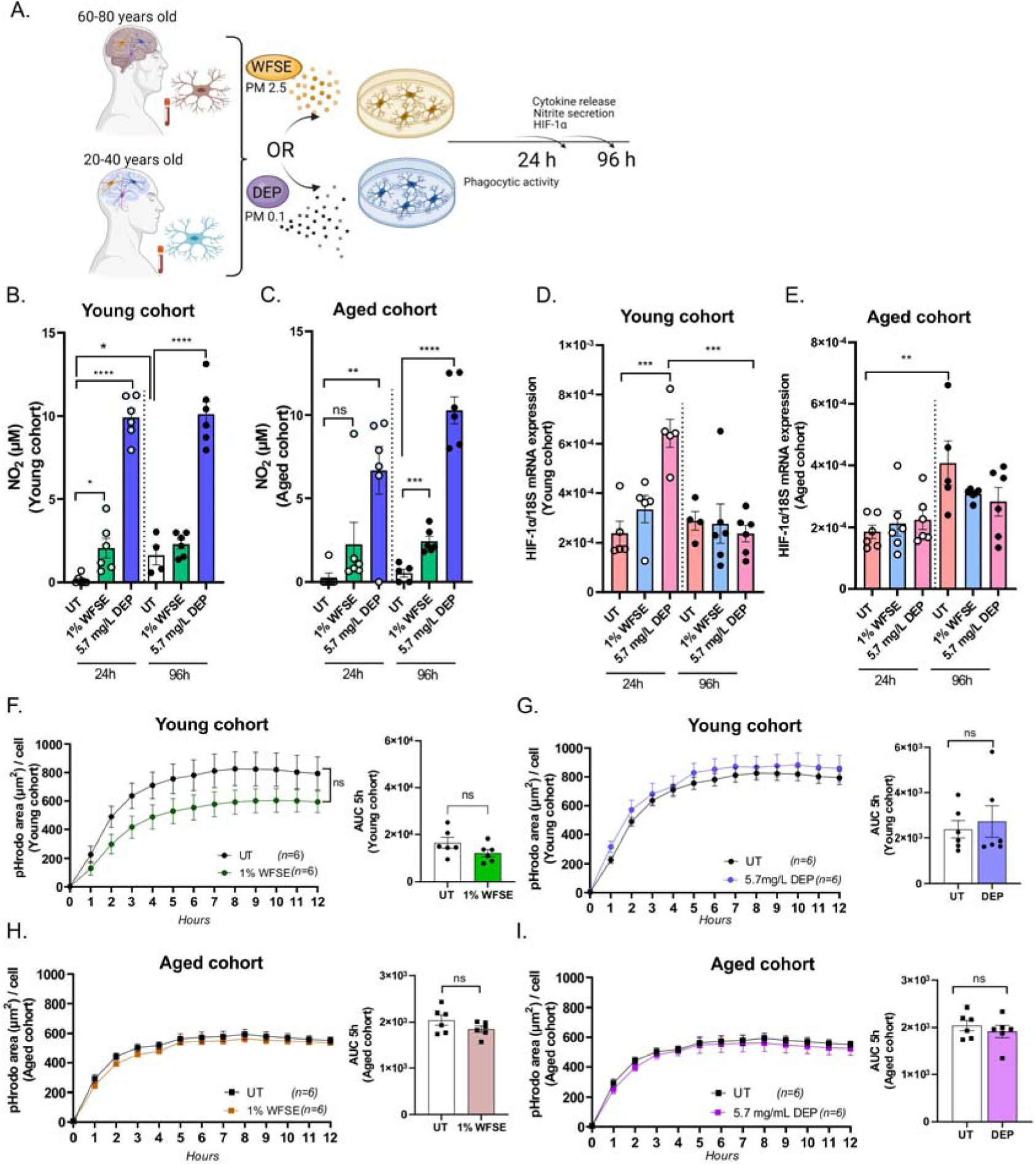
WFSE and DEP induce nitrosative stress without altering phagocytic capability in MDMi. **(A)** Young and aged MDMi exposed to either 1% WFSE or 5.7 mg/L DEP, and MDMi were harvested at 24 hr and/or 96 hr to assess phagocytic activity, nitrosative stress, and transcriptional regulation of HIF-1. **(B)** Nitrosative stress levels in young and **(C)** aged MDMi were measured following exposure to 1% WFSE or 5.7 mg/L DEP at 24 hr and/or 96 hr. **(D)** HIF-1α mRNA expression was determined in young MDMi and **(E)** aged MDMi after treatment with 1% WFSE or 5.7 mg/L DEP at 24 hr and 96 hr. **(F-G)** Phagocytic activity in young MDMi and **(H-I)** aged MDMi was evaluated by tracking the uptake of pH-sensitive fluorescent *E. coli*-coated BioParticles over time (n = 6). Data presented as mean ± SEM, with data points representing biological replicates (donors). Statistical analysis between two groups was performed using Student’s *t* test and between multiple groups using one-way ANOVA. Statistical significance indicated as follows: *\**p < 0.05, **p < 0.01, ***p < 0.001, ****p < 0.0001.

Elevated nitrate secretion, a marker of NO production was observed in young MDMi treated with 1% WFSE at 24 hr against untreated control (p = 0.021, Fig. 2B), with these levels remaining consistent through to 96 hr. In aged MDMi, total NO levels showed a slight elevation from 24 to 96 hr, with notable variability among patients in 24 hr (Fig. 2C). Comparative analysis between the young and aged MDMi cohorts showed similar NO levels at both 24 hr and 96 hr, indicating no significant effect of age (**Fig. S2A-C)**.

Exposure to 5.7 mg/L DEP induced a significant increase in NO levels in both young and aged MDMi at 24 hr and 96 hr against untreated control, indicating that DEP can induce substantial nitrosative stress within 24 hr, which persists at 96 hr, regardless of age. Moreover, young MDMi appear to respond more robustly than aged MDMi, as NO levels at 24 hr were similar to those at 96 hr (24 hr vs 96 hr, p = 0.84, Fig. 2B). In contrast, a trending increase in NO levels was observed in aged MDMi treated with DEP (24 hr vs 96 hr, p = 0.053, Fig. 2C). A significant age effect was observed at 24 hr, with increased NO levels in young compared to aged MDMi, which normalised at 96 hr (**Fig. S2A-C**), supporting the robust NO response in young MDMi. Collectively, DEP exposure induced stronger NO responses in both young and aged MDMi than WFSE, with aged cells showing a delayed response in both air pollutants.

NO can induce hypoxia-inducible factor 1 alpha (HIF-1α), which accumulates under hypoxic and inflammatory conditions associated with sustained NO production [55]. Supporting this, previous work showed NO secretion in response to air pollutants, such as PM 2.5, increased HIF-1α expression in endothelial cells (HUVEC) [56]. To address the possibility of HIF-1α accumulation, total RNA was extracted from young and aged MDMi exposed to 1% WFSE or 5.7 mg/L DEP for 24 hr and 96 hr, and the level of mRNA expression was examined by RT-qPCR. We found that HIF- 1α expression significantly increased in young MDMi exposed to 5.7 mg/L DEP at 24 hr compared to untreated control (p = 0.0005), returning to baseline by 96 hr (p = 0.0003, Fig. 2D). No differences in HIF-1α expression were observed in aged MDMi exposed to DEP or WFSE compared to untreated control. However, an increase in basal HIF-1α gene expression was noted in vehicle-treated aged MDMi from 24 hr to 96 hr (p = 0.039), which was not seen in young MDMi (Fig. 2E**, Fig. S2D**). A comparative analysis between young and aged MDMi cohorts revealed a significant age effect at 24 hr, with higher HIF-1α levels in young MDMi compared to aged MDMi, which normalised by 96 hr (**Fig. S2D-F**). This finding highlights that young MDMi may respond more dynamically to environmental stressors, while aged MDMi might exhibit delayed or dampened responses.

Phagocytosis is a key protective function of microglia that can be impacted by toxic stimuli such as air pollutants. Further, existing studies using immortalised macrophage (THP-1) and microglia (SV40) cultures have indicated that exposure to air pollution can adversely impact phagocytosis [20, 48, 57]. However, the phagocytic response of MDMi to air pollution remains unknown. To test this, young and aged MDMi were treated with 1% WFSE or 5.7 mg/L of DEP for 1 hr before the addition of pH-sensitive fluorescent *E. coli-* coated BioParticles (pHrodo BioParticles). Live imaging was performed to determine uptake of pHrodo BioParticles hourly over 12 hrs. Area under the curve (AUC) indicated a moderate, but non-significant, decrease (p = 0.138) in phagocytosis in young MDMi treated with 1% WFSE (Fig. 2F), while no differences were observed in 5.7 mg/L DEP-treated young MDMi (p = 0.67, Fig. 2G) compared to control. Similar uptake of pHrodobeads was observed in aged MDMi treated with 1% WFSE (p = 0.163, Fig. 2H) and 5.7 mg/L DEP (p = 0.48, Fig. 2I) compared to control. Additionally, we noted a moderate decrease in phagocytic activity in aged MDMi compared to young MDMi in control samples (treated with pHrodobeads only), suggesting that ageing itself may influence microglial phagocytosis (**Fig. S2G**).

### Short-and long-term exposure to air pollutants increases inflammatory response in MDMi

To investigate the effects of short- and long-term air pollution exposure, both young and aged MDMi were treated with 1% WFSE or 5.7 mg/L DEP, and cell media were sampled at multiple time points (2 hr, 24 hr, 48 hr, and 96 hr; Fig. 3A). Cytokine secretion was profiled using a 13-plex human inflammation panel, excluding IL-1β, IL-12p70, and IL-17A due to levels below the detection limit.

**Figure 3.**
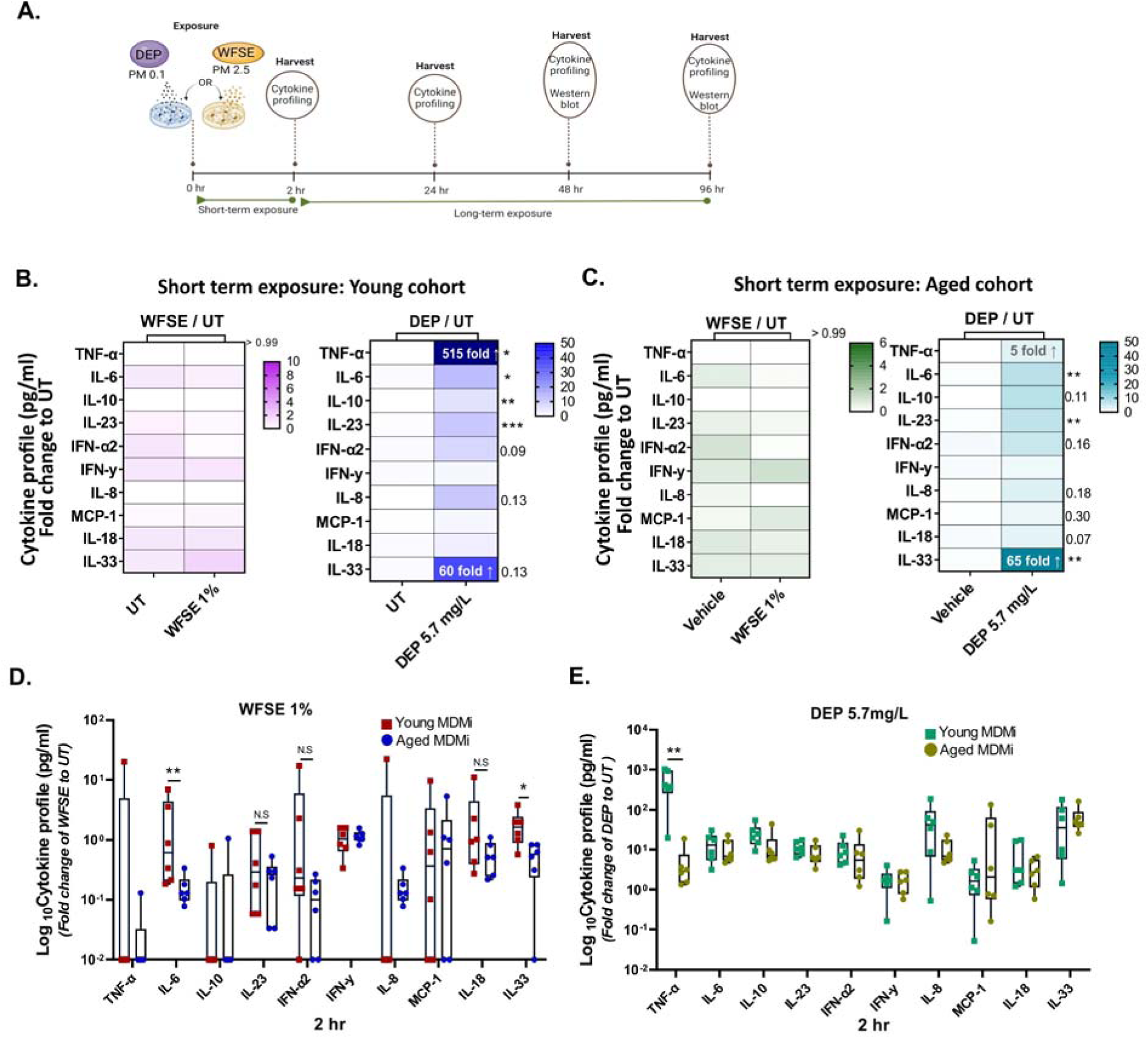
Rapid and pronounced cytokine response to short-term DEP exposure compared to WFSE in both MDMi cohorts. **(A)** Schematic of exposure timeline, showing young and aged MDMi exposure to either 1% WFSE or 5.7 mg/L DEP for short-term (2 hr) and long-term (24 hr, 48 hr, and 96 hr) exposures. **(B)** Cytokine profiling in young MDMi exposed to 1% WFSE (left) or 5.7 mg/L DEP (right), normalised to untreated controls. **(C)** Cytokine profiling in aged MDMi exposed to 1% WFSE (left) or 5.7 mg/L DEP (right), normalised to untreated controls after 2 hr. **(D)** Comparative cytokine profiling between young and aged MDMi exposed to 1% WFSE, and **(E)** between young and aged MDMi exposed to 5.7 mg/L DEP. Data presented as mean ± SD, with data points representing biological replicates (donors). Statistical analysis between two groups was performed using Student’s *t* test and between multiple groups using one-way ANOVA or two-way ANOVA Statistical significance indicated as follows: *p < 0.05, **p < 0.01, ***p < 0.001, ****p < 0.0001.

### DEP induces rapid cytokine responses compared to WFSE in young and aged MDMi

In young MDMi exposed to 5.7 mg/L DEP, significantly elevated levels of TNF-α, IL-6, IL-10, and IL-23 were observed as early as 2 hr, while cells treated with 1% WFSE showed no significant changes compared to vehicle controls at this time point (Fig. 3B**, Fig. S3A**). Similarly, aged MDMi exposed to 5.7 mg/L DEP displayed increased IL-6, IL-23, and IL-33, with no significant changes detected in the 1% WFSE-treated cells compared to vehicle controls (Fig. 3C**, Fig. S3B**). These findings indicate that DEP induces a pronounced and rapid inflammatory response in both cohorts at early exposure (2 hr). Notable cytokine increases included a 515-fold rise in TNF-α in young MDMi (p = 0.013) and a 65-fold rise in IL-33 in aged MDMi (p = 0.008) compared to untreated controls, highlighting distinct cytokine profiles between young and aged MDMi under early DEP exposure (Fig. 3B**-C****, Fig. S3C-D**).

Additionally, comparative analysis of young and aged MDMi at 2hr revealed a significant increase in IL-6 and IL-33 responses following exposure to 1% WFSE (Fig. 3D). Similarly, a higher TNF-α response was seen in young MDMi compared to aged MDMi upon exposure to 5.7 mg/L DEP (Fig. 3E). These findings suggest that the diminished IL-6 and IL-33 responses to early WFSE exposure, as well as the reduced TNF-α response to DEP, may be associated with ageing and impair aged microglia to mount effective responses to toxic pollutants.

### Chronic WFSE and DEP exposure drives inflammatory responses in aged MDMi

Extended exposure to air pollution is a well-established health hazard with detrimental long-term effects. However, the specific inflammatory responses driven by microglia, particularly in aged individuals remain poorly characterised. To address this, we evaluated the inflammatory profiles of young and aged MDMi following prolonged exposure to 1% WFSE and 5.7 mg/L DEP. Supernatants were collected at 2 hr, 24 hr, 48 hr, and 96 hr for cytokine profiling to capture dynamic inflammatory responses.

### Prolonged WFSE exposure increase inflammatory responses in aged MDMi

Prolonged exposure to 1% WFSE induced a time-dependent increase in cytokine secretion, with aged MDMi exhibiting a markedly amplified inflammatory response compared to their younger counterparts. By 96 hr, aged MDMi demonstrated significant upregulation in 8 out of 10 cytokines, including TNF-α (31.5-fold, p = 0.0089), IL-33 (75.4-fold, p < 0.0001), IFN-γ, IL-23, IL-10, IL-18 (all p < 0.0001), and IFN-α2 (p = 0.005) against untreated control (Fig. 4A**-B****, Fig. S4A-B**). In contrast, young MDMi showed significant elevation in only 3 cytokines under the same conditions (Fig. 4A**, Fig. S4A**). In both cohorts, MCP-1, IL-6, and IL-8 displayed gradual increases over time but remained comparable to untreated controls (**Fig. S4A-B**).

**Figure 4.**
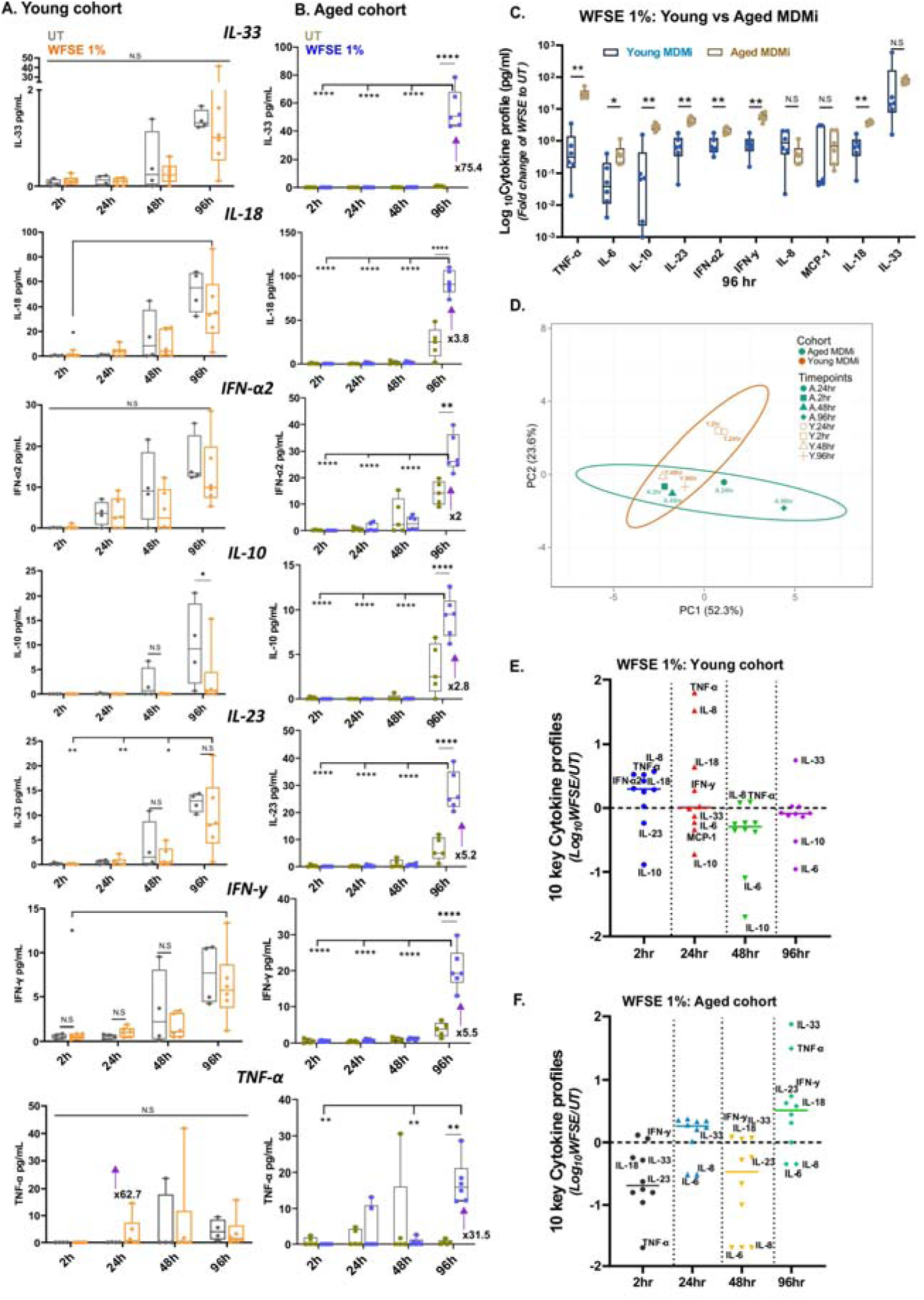
Long-term exposure to WFSE 1% reveals enhanced inflammatory response in aged MDMi. **(A)** Longitudinal time-course analysis of young or **(B)** aged MDMi treated with WFSE 1% or untreated control shows a time-dependent response of cytokines. **(C)** Comparative analysis of young and aged MDMi at 96 hr. **(D)** Principal-component analysis (PCA) was performed on log-transformed cytokine profiles across 2 hr, 24 hr, 48 hr, and 96 hr. PCA reveals distinct clustering patterns between young and aged MDMi discriminated by time-points. Overlapping time points indicate similar responses. Singular value decomposition (SVD) with imputation is used to calculate principal components. X and Y axis show PC1 and PC2 that explain 52.3% and 23.6% of the total variance, respectively. **(E)** The graph highlights 10 key cytokines from (D) that drive PCA separation in young MDMi across all time points. TNF-α and IL-8 exhibited pronounced increases at 24 hr, while IL-33 showed the highest increase at 96 hr in cells exposed to 1% WFSE. **(F)** The graph depicts 10 key cytokines from (D) driving PCA separation in aged MDMi across all time points. Elevated levels of IL-33 and TNF-α were observed at 96 hr in cells exposed to 1% WFSE. Fold-change values are normalised to the untreated control at each specific time point then log-transformed. Data are presented as mean ± SD, with data points representing biological replicates (donors; young, n = 4-6; aged, n = 5-6). Statistical significance determined using Student’s *t*-test for two-group comparisons and one-way or two-way ANOVA for multiple groups. *p < 0.05, **p < 0.01, ***p < 0.001, ****p < 0.0001.

Age-related differences in cytokine dynamics were particularly pronounced. Aged MDMi exhibited elevated IFN-γ levels at 48 hr (p = 0.0087), which persisted through 96 hr (p = 0.0022) compared to young MDMi (**Fig. S4D**). By 96 hr, aged MDMi showed significant upregulation of TNF-α (p = 0.0022), IL-6 (p = 0.0260), IL-10 (p = 0.0022), IL-23 (p = 0.0022), IFN-α2 (p = 0.0043), and IL-18 (p = 0.0022) relative to young MDMi, while MCP-1, IL-8, and IL-33 levels remained similar across both cohorts (Fig. 4C).

Principal component analysis (PCA) was used to discriminate study groups by exposure time, with PC1 and PC2 explaining 52.3% and 23.6% of the total variance. Analysis of 10 key log-transformed cytokines across all time points revealed cohort-specific shifts, with young MDMi showing the most significant changes at 2 hr and 24 hr, driven primarily by elevated TNF-α and IL-8 levels (Fig. 4D**-E****, Fig. S4E**). In contrast, aged MDMi exhibited their most pronounced changes at 96 hr followed by 24 hr, characterised by elevated IL-33 and TNF-α levels (Fig. 4D**-F****, Fig. S4E**).

These findings demonstrate that prolonged WFSE exposure triggers an amplified and sustained inflammatory response in aged MDMi compared to young MDMi, underscoring age-related susceptibility to air pollution-induced microglial activation. Such differences highlight the potential for heightened vulnerability in aged individuals to chronic environmental exposures.

### DEP exposure reveals delayed and amplified cytokine profiles in aged MDMi compared to young

Extended exposure to DEP 5.7 mg/L induced a time-dependent cytokine increase from 2 hr to 96 hr in both young and aged MDMi. Young MDMi displayed the most significant increases in TNF-α (563.1-fold change at 24 hr, p =0.01) and IL-33 (70.1-fold change at 24 hr, p = 0.005) after DEP exposure compared to untreated control (Fig. 5A**, Fig. S5A**). Similarly, aged MDMi exhibited significant increases in IL-10 (134.2-fold, p = 0.004), IL-33 (131.3-fold, p = 0.004), and IL-23 (18-fold, p = 0.004) at 24 hr (Fig. 5B**, Fig. S5B**). Most cytokine levels normalised between 48 hr to 96 hr, except for IL-33, which remained significantly elevated in both young (28-fold, p < 0.0001) and aged MDMi (42-fold, p < 0.0001) at 96 hr. This sustained IL-33 elevation suggests a specific and prolonged response to DEP exposure.

**Figure 5.**
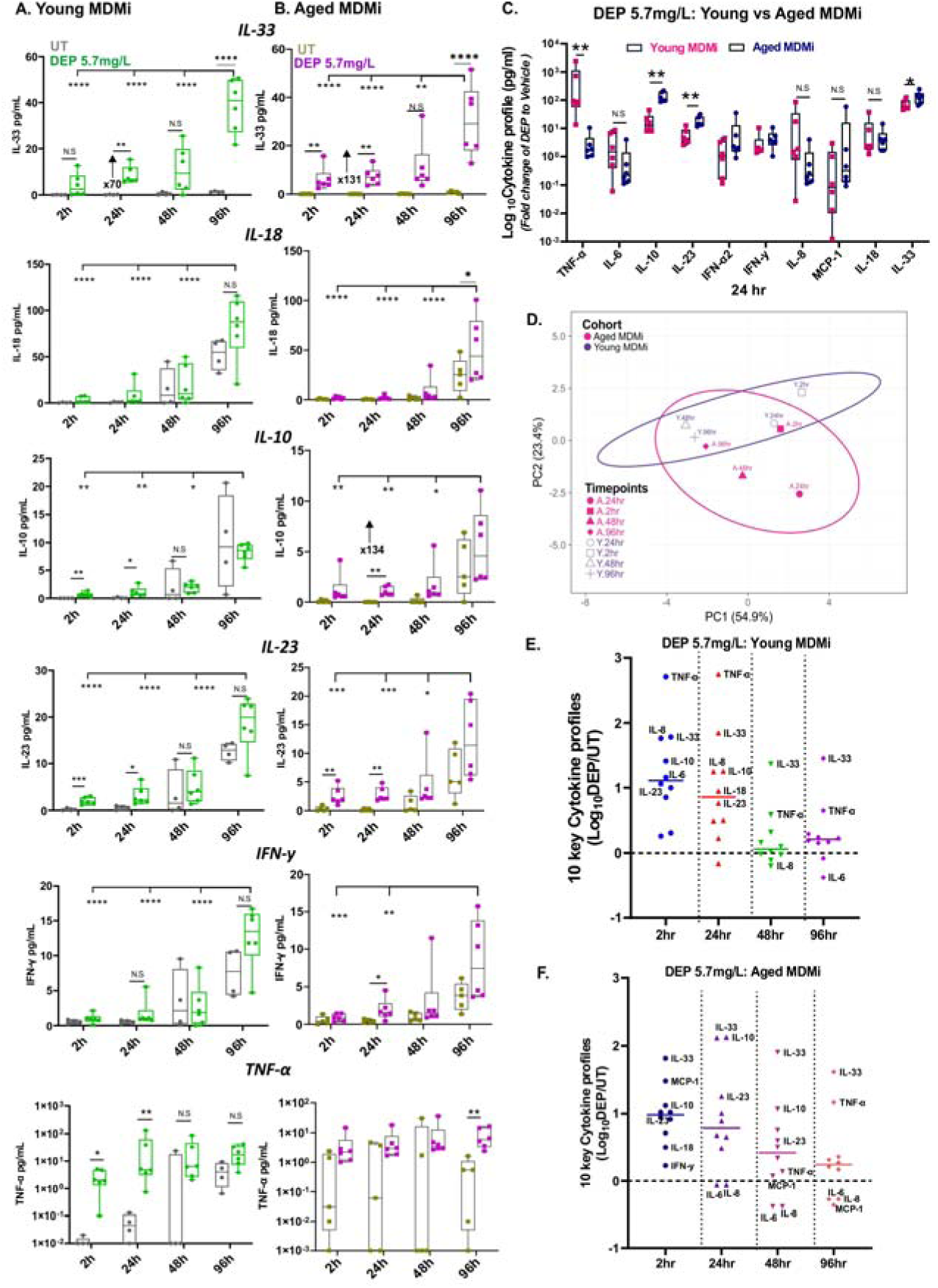
Aged MDMi exhibit peak cytokine response at 24 hr following DEP exposure. **(A-B)** Longitudinal time-course analysis of cytokine levels in young and **(B)** aged MDMi treated with DEP 5.7 mg/L or untreated control, showing a time-dependent increase in cytokine levels. Most cytokines normalised by 48 hr, except for IL-33, which remained elevated at 96 hr in both young and aged MDMi. **(C)** Comparative cytokine response between aged and young MDMi at 24 hr following DEP exposure. Aged MDMi exhibit consistently higher levels of IL-10, IL-23, and IL-33 at 24 hr and show prolonged elevation of IL-10 and IL-33 compared to young MDMi. **(D)** PCA of log-transformed cytokine profiles across 2 hr, 24 hr, 48 hr, and 96 hr. In young MDMi, major shifts are observed at 2 hr while in aged MDMi changes occur at 24 hr. Overlapping time points indicate similar responses. SVD with imputation is used to calculate principal components. X and Y axis show PC1 and PC2 that explain 54.9% and 23.4% of the total variance, respectively. **(E)** The graph highlights 10 key cytokines from (D) driving PCA separation in young MDMi across all time points. TNF-α and IL-33 exhibited pronounced increases at 2 hr and 24 hr following DEP exposure. **(F)** The graph depicts 10 key cytokines from (D) driving PCA separation in aged MDMi across all time points, with significant elevations in IL-33, IL-10, and IL-23 at 24 hr following DEP exposure. Fold-change values are normalised to the vehicle control at each specific time point. Data presented as mean ± SD, with data points representing biological replicates (donors; young, n = 4-6; aged, n = 5-6). Statistical significance determined using Student’s *t*-test for two-group comparisons and one-way or two-way ANOVA for multiple groups. Significance levels indicated as follows: *\**p < 0.05, **p < 0.01, ***p < 0.001, ****p < 0.0001.

Comparative analysis between young and aged MDMi revealed significant cohort-specific differences in cytokine responses. TNF-α levels were lower in aged MDMi at 24 hr compared to young MDMi (p = 0.002) but increased significantly by 96 hr (p = 0.04) (Fig. 5C**, Fig. S5C-D**). Additionally, aged MDMi displayed consistent elevations in IL-10 (p = 0.002), IL-23 (p = 0.004), and IL-33 (p = 0.04) at 24 hr, with IL-10 (p = 0.002) and IL-33 (p = 0.037) remaining elevated at 48 hr compared to young MDMi (**Fig. S5C-D**). These findings indicate that aged MDMi exhibit a more prolonged and exacerbated cytokine response to DEP exposure than young MDMi.

To further evaluate the dynamics of DEP-induced cytokine secretion, a PCA of 10 cytokines was performed across time points. PCA revealed cohort-specific patterns, with young MDMi showing pronounced cytokine shifts at 2 hr, driven by elevated TNF-α, IL-8, and IL-33 levels (Fig. 5D**-E****, Fig. S5E**). In contrast, aged MDMi displayed significant increases in IL-10 and IL-33 at 24 hr, highlighting a delayed (compared to young) but robust response (compared to WFSE) (Fig. 5D**-F****, Fig. S5E**).

Overall, these results demonstrate distinct inflammatory dynamics in response to DEP exposure, with aged MDMi exhibiting a more prolonged and amplified cytokine response compared to young MDMi. While DEP exposure induced early cytokine shifts in young MDMi (2 hr), aged MDMi displayed a delayed (24 hr) yet sustained inflammatory response, highlighting age-related differences in susceptibility to air pollution-induced microglial activation. Notably, considerable intra-individual variability was observed within both age cohorts, with some donors showing markedly different responses to DEP exposure compared to cohort averages. This variability likely reflects individual-specific factors, including genetic background or baseline immune states, as well as potential cohort effects, including differences in environmental exposure history. These findings highlight the complexity of the inflammatory response to air pollution and underscore the importance of accounting for both individual and cohort-level variability in future studies.

### WFSE and DEP induced antioxidant gene HO-1 expression and the phosphorylation of ERK, p38 MAPK and NF-**κ**B p65 nuclear translocation

Air pollutants promotes oxidative stress and inflammation, contributing to neurotoxicity and immune dysregulation in various cell types, including those in the brain [58, 59]. To investigate air pollutant-induced inflammation, we examined HO-1 activation, a key antioxidant gene involved in oxidative stress responses across various cell types, including macrophages [60, 61].

Young MDMi were exposed to 1% WFSE or 5.7 mg/L DEP for 24 hr and 96 hr to assess HO-1 protein expression by immunoblotting. At 24 hr, expression of HO-1 increased significantly following exposure to either WFSE or DEP compared to controls (p < 0.0001). By 96 hr, DEP and WFSE-induced HO-1 expression returned to baseline compared to control (Fig. 6A**-B**). These findings suggest that air pollutants trigger acute antioxidant responses, HO-1, in MDMi that resolves by 96 hr.

**Figure 6.**
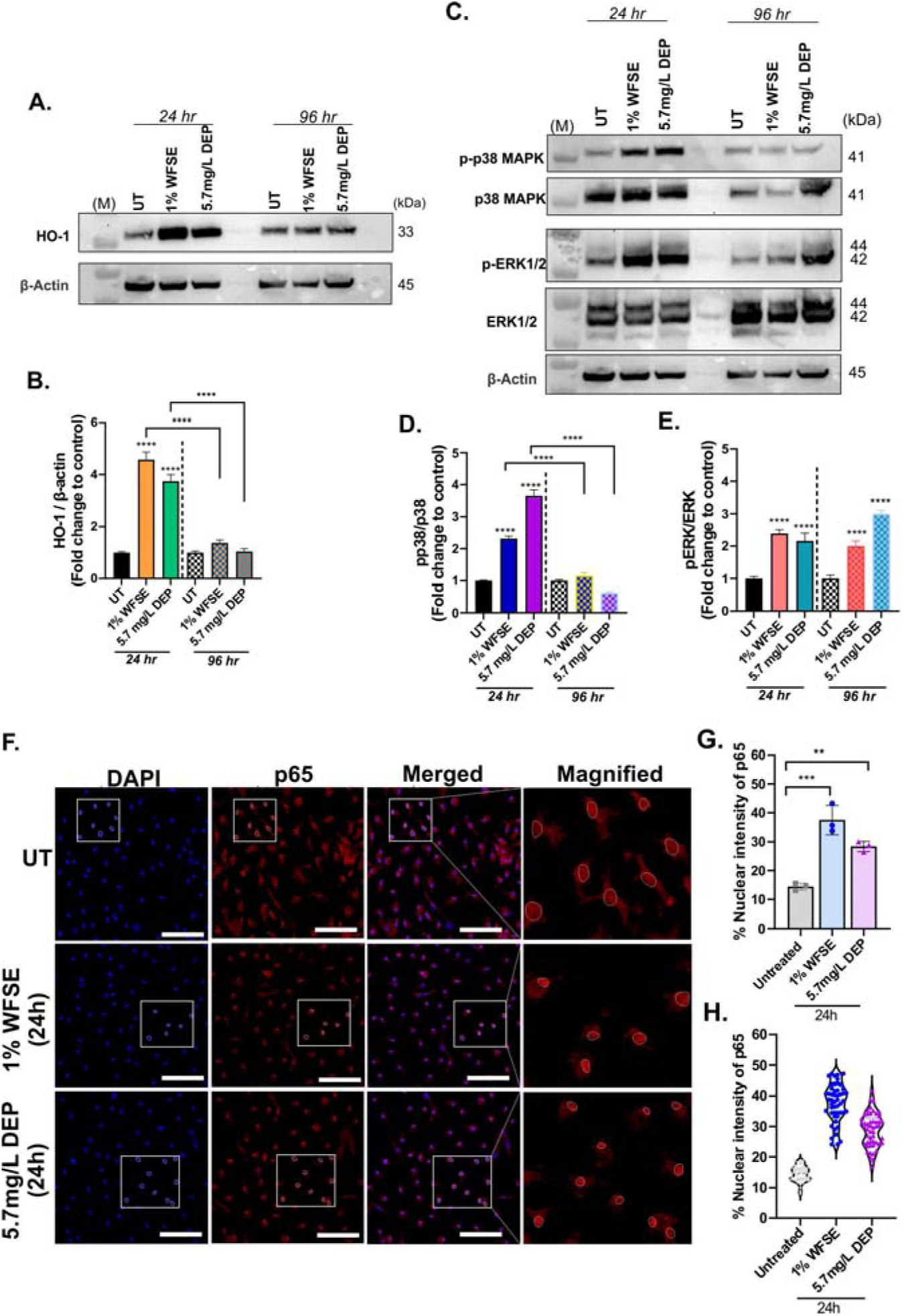
WFSE and DEP induced HO-1 expression, phosphorylation of p-38, ERK1/2 and Activation of the NF-κB pathway in young MDMi upon air pollutant exposure. **(A)** Representative western blot images showing HO-1 protein expression in young MDMi exposed to 1% WFSE or 5.7 mg/L DEP for 24 hr and 96 hr. **(B)** Quantification of HO-1 expression at 24 hr and 96 hr. At 24 hr, both WFSE and DEP significantly increased HO-1 levels compared to untreated controls **(C)** Representative western blot images showing phosphorylation of p38 (p-p38), total p38, p-ERK1/2, and total ERK1/2 protein expression in young MDMi exposed to 1% WFSE or 5.7 mg/L DEP for 24 hr and 96 hr. **(D)** Quantification of p-p38 levels normalised to total p38. **(E)** Quantification of p-ERK1/2 levels normalised to total ERK1/2. All proteins were normalised to β-actin before fold-change was performed. **(F)** Representative immunofluorescence images showing the localisation of NF-κB p65 in young MDMi at 24 hr following exposure to WFSE or DEP. Dotted circles in the panels indicate nuclear localisation as marked by DAPI staining. **(G-H)** Quantification of cells that has nuclear translocation of p65, total n = 50 cells analysed from n = 3 donors. **(H)** Shows the level of nuclear p65 intensity in individual cells across WFSE and DEP exposure. Data presented as mean ± SEM, n = 3. Scale bars represent 50 µm. UT= untreated. Significance levels indicated as follows: **p < 0.01, ***p < 0.001, ****p < 0.0001.

To further elucidate the signalling pathways involved in the observed air pollutant-induced inflammation and HO-1 expression in MDMi, we focused on the MAPK pathways and key pro-inflammatory transcription factors such as NF-κB p65, both of which can be directly activated by air pollutant-induced inflammatory responses in various cell types, including macrophages [59, 62–65].

We assessed the p38 and ERK MAPK signalling pathways in MDMi exposed to WFSE 1% or DEP 5.7 mg/L for 24 hr and 96 hr. Both p38 and ERK activation peaked at 24 hr in WFSE- and DEP-treated MDMi (Fig. 6C**-E**). By 96 hr, phosphorylated-p38 level returned to baseline in both treatments, while ERK activation sustained up to 96 hr in both WFSE- and DEP-treated MDMi compared to controls. These results suggest that WFSE and DEP exposure induce initial activation of both p38 and ERK MAPK pathways at 24 hr, but only ERK shows sustained activation. Taken together, these results show that both air pollutants activate the p38 and ERK pathways in a time-dependent manner and that the resolution of p38 and prolonged ERK signalling may likely indicate distinct inflammatory/stress adaptation mechanisms.

To investigate the activation of the air pollutant-induced NF-κB pathway, young MDMi were exposed with WFSE or DEP for varying durations. Immunofluorescence staining revealed significant nuclear translocation of NF-κB p65 at 24 hr in both WFSE- and DEP-treated MDMi, indicating activation of the NF-κB pathway. No significant changes in p65 localisation were observed at earlier time points (15 min, 30 min and 1 hr, **Fig. S6**), suggesting a delayed response to air pollutant exposure. These findings highlight the time-dependent NF-κB activation, with nuclear translocation occurring within 24 hr following air pollutant exposure (Fig. 6F**-H**).

## Discussion

Air pollutants exert their effects predominantly through local exposure via the olfactory system, which serves as a primary entry route for airborne toxins into the brain [24]. Evidence from studies on dogs in Mexico City revealed not only nasal epithelial damage but also mononuclear cell infiltration, increased neuroinflammatory markers, ultrafine particle-like material in red blood cells, and amyloid-beta deposits in the brain [66]. Similar observations have been made in experimental models, where prolonged exposure to wildfire smoke and DEP induced neuroinflammation characterised by microglial activation and the infiltration of peripheral immune cells into the CNS [67, 68]. These studies highlight the capacity of WFSE and DEP to promote neuroinflammation, in part by facilitating the infiltration of inflammatory monocytes into the brain, potentially accelerating neurological ageing and pathology.

Despite these advances, the fate of air pollutant-induced infiltrating monocytes within the brain remains unclear. Addressing this gap, our study utilised a monocyte-derived microglia model to explore air pollution exposure’s impact on brain immune responses and its role in ageing and neurodegenerative processes. Our results revealed that DEP and WFSE elicit distinct time-dependent inflammatory responses in MDMi from young and aged cohorts. Specifically, DEP triggered acute increases in TNF-α, IL-6, IL-23, and IL-33 within 2 hr, aligning with prior reports of DEP-driven microglial activation [67] and IL-33 upregulation in epithelial cells [69, 70] and macrophages exposed to PM2.5 [71]. Notably, IL-23, though less characterised in the context of air pollution, likely plays a dual role in innate and adaptive immunity through recruitment of inflammatory cells [72]. In contrast, WFSE induced a delayed yet sustained inflammatory response, particularly in aged MDMi, with TNF-α and IL-33 levels persisting at 96 hr. This finding is consistent with previous studies reporting prolonged neuroinflammation following wildfire-derived particulate exposure [68, 73]. Together, these findings support the hypothesis that both WFSE and DEP drives microglial activation contributing to neurological ageing and pathology.

Age emerged as a critical factor modulating these responses, with MDMi from older people exhibiting amplified and prolonged inflammatory profiles to WFSE compared to DEP, including upregulation of 7 out of 10 cytokines studied (e.g., TNF-α, IL-6, IL-10, IFN-γ, and IL-33) relative to younger cells. This heightened response likely reflects age-related vulnerabilities, such as increased DNA damage [74, 75], which may contribute to the elevated cytokine levels observed here and in alveolar macrophages and epithelial cells exposed to PM2.5 pollutants [76]. Age-associated deficits in immunometabolism may further impair microglial regulation and homeostasis [77–79] including NF-κB activation and cytokine induction [80]. Interestingly, these metabolic alterations were more pronounced in microglia derived from females in Alzheimer’s disease contexts [81, 82]. Similar findings have been observed in aged mice exposed to wildfire smoke, which demonstrated prolonged inflammation, neurological deficits, and metabolic changes linked to aging [83]. Conversely, aged MDMi displayed diminished early cytokine responses, such as IL-6 and IL-33 to WFSE and TNF-α to DEP, potentially due to impaired early cytokine signalling pathways. IL-33, in particular, plays a crucial role in mitigating β-amyloid accumulation, as evidenced by APP/PS1 mouse models where IL-33 deficiency exacerbates Alzheimer’s disease pathology [84].

The distinct inflammatory profiles elicited by DEP and WFSE likely stem from their differing physicochemical properties, where a potentially smaller particle size in DEP could have a larger impact on toxicity and inflammatory outcomes [21, 85–87]. DEP’s ultrafine particles, for example, are likely more adept at penetrating the CNS, triggering acute inflammation and nitrosative stress [86, 88], while WFSE, derived from burned vegetation [48], comprises a complex chemical mixture, which sustains inflammation through prolonged microglial activation. DEP exposure resulted in robust NO production, particularly in younger MDMi, consistent with heightened inducible NO synthase activity observed in post-mortem brains exposed to air pollution [16] and in murine microglial models exposed to PM2.5 [52]. In contrast, aged MDMi exhibited reduced and delayed NO responses compared to young MDMi at 24 hr exposed to DEP (**Fig S2A, 2C**), reflecting age-associated impairments in inflammatory resilience [29, 54]. These findings potentially highlight aging microglia’s diminished ability to overcome environmental stressors, potentially exacerbating cell stress and contributing to neurodegenerative vulnerability in elderly populations.

Oxidative stress is a central mechanism underlying air pollutant-induced inflammation, driven by an imbalance between reactive oxygen species production and the capacity of antioxidant defence systems. HO-1, a highly inducible anti-inflammatory enzyme, plays a critical role in mitigating oxidative stress and inflammation during such challenges. Our study demonstrates that exposure to both WFSE and DEP significantly upregulates HO-1 expression by 24 hr, with levels normalising by 96 hr (Fig. 6B), indicating a resolution of oxidative stress. Additionally, exposure to both DEP and WFSE altered HIF-1α expression, with young MDMi showing significantly increased HIF-1α in response to DEP (24 hr), while aged MDMi exhibited elevated basal HIF-1α regardless of exposure (**Fig. S2F**). This suggests a chronic low-level inflammatory state in aged microglia that may impair their adaptive capacity to subsequent inflammatory insults, exacerbating neurodegenerative risk.

To explore the signalling pathways mediating air pollutant-induced inflammatory responses, we examined MAPK and NF-κB activation. Both DEP and WFSE exposures activated p38 MAPK, ERK1/2, and NF-κB p65 by 24 hr. While p38 activation resolved by 96 hr, ERK activation persisted, reflecting distinct roles for these pathways in stress and proliferation responses respectively. p38 activation aligns with its established function in cellular stress responses, particularly in acute inflammation, oxidative stress, and environmental stimuli. The transient nature of p38 likely reflects a rapid but temporary cellular adaptation to restore homeostasis. In contrast, sustained ERK activation indicates prolonged pro-inflammatory signalling, which may contribute to the persistence of inflammatory cascades. These findings are consistent with prior studies showing similar ERK and p38 activation in DEP-treated human bronchial epithelial cells [61]. The nuclear translocation of NF-κB p65 observed following DEP and WFSE treatment, suggest its critical role in driving the cytokine release observed in this study, and are consistent with previous reports of NF-κB activation in DEP-exposed bronchial epithelial cells [89, 90], in mice exposed to PM [91, 92], BV2 murine microglia lines [93] and post-mortem brain tissues from regions with high air pollution exposure [16]. To the best of our knowledge, NF-κB p65 activation in microglia exposed to WFSE has not been previously reported. Here, we provide the first evidence of NF-κB activation in human-derived MDMi in response to both DEP and WFSE.

Additionally, our study revealed distinct impacts of air pollutants on microglial phagocytic function. WFSE exposure moderately reduced phagocytic activity in young MDMi, while DEP did not impair phagocytosis in this model. This contrasts with rat microglia studies linking DEP to increased superoxide as an indirect presence of microglial phagocytosis [25]. WFSE exposure, on the other hand and cigarette smoke extract exposed macrophages (THP-1 and monocyte-derived) show decreased phagocytosis that is coupled with altered phagocytic receptors [48, 94]. This disparity in results may be due to the type of fuel, engine operating conditions (for DEP) and different concentrations, exposure (for WFSE) and analysis methods in microglial phagocytosis, highlighting the need for further research. Additionally, aged MDMi displayed moderately reduced baseline phagocytic activity (**Fig. S2G**), further implicating aging in compromised microglial function and heightened susceptibility to environmental pollutants.

Together, these findings demonstrate the distinct age- and pollutant-specific inflammatory, oxidative, and functional responses in MDMi. DEP elicits acute, transient effects driven by its ultrafine nature, while WFSE induces sustained inflammatory and oxidative stress, particularly in aged microglia. These differential responses underscore the vulnerability of aging microglia to environmental pollutants and their potential role in exacerbating age-related neurodegeneration.

### Limitations of the study

In addition to PM, toxic gases such as carbon monoxide and ozone (OL) can further exacerbate neurological risk such as the accumulation of amyloid-beta in aged individuals [95]. However, our current methods used for air pollutant collection does not account for the full profile of associated gaseous pollutants. Furthermore, the chemical composition of PM is highly heterogeneous, containing diverse trace metals that can influence bioaccumulation and disrupt the redox imbalance in cells leading to oxidative stress [96]. This heterogeneity is particularly pronounced in wildfire smoke, where the composition varies depending on the materials being burned. The extract used in this study specifically represents smoke derived from Australian plant matter and excludes toxic compounds that could be released when bushfires burn housing or infrastructure alongside vegetation. Understanding the chemical composition of PM is crucial for accurately predicting its inflammatory and oxidative effects, enabling the development of targeted strategies to mitigate the health impacts of PM exposure based on its source, composition, and exposure duration.

## Conclusion

We show that both WFSE and DEP trigger microglial activation, as evidenced by the up-regulation of pro-inflammatory mediators, cytokines and chemokines, and nitrosative stress in both young and aged MDMi. DEP induced a stronger inflammatory and nitrosative response in short-term exposure (acute) while WFSE induced a stronger inflammatory response in long-term exposure (chronic), particularly in aged microglia. These findings suggest that the type and duration of air pollution exposure influence the inflammatory response of microglia in different ways. Additionally, our results also show that ageing microglia may be more vulnerable to prolonged exposure to WFSE.

We also identified that both NF-κB p65 and MAPK signalling pathway play a role in the up-regulation of pro-inflammatory cytokines in microglia following WFSE and DEP exposure, as evidenced by nuclear translocation of NF-κB and the phosphorylation of p-38 and ERK. These results suggest a potential mechanism and target pathway involved in microglia response towards air pollutants in brain health (Fig. 7). Further research in this area may contribute to the development of strategies to mitigate the adverse effects of air pollution on brain health, particularly in vulnerable populations such as the elderly.

**Figure 7.**
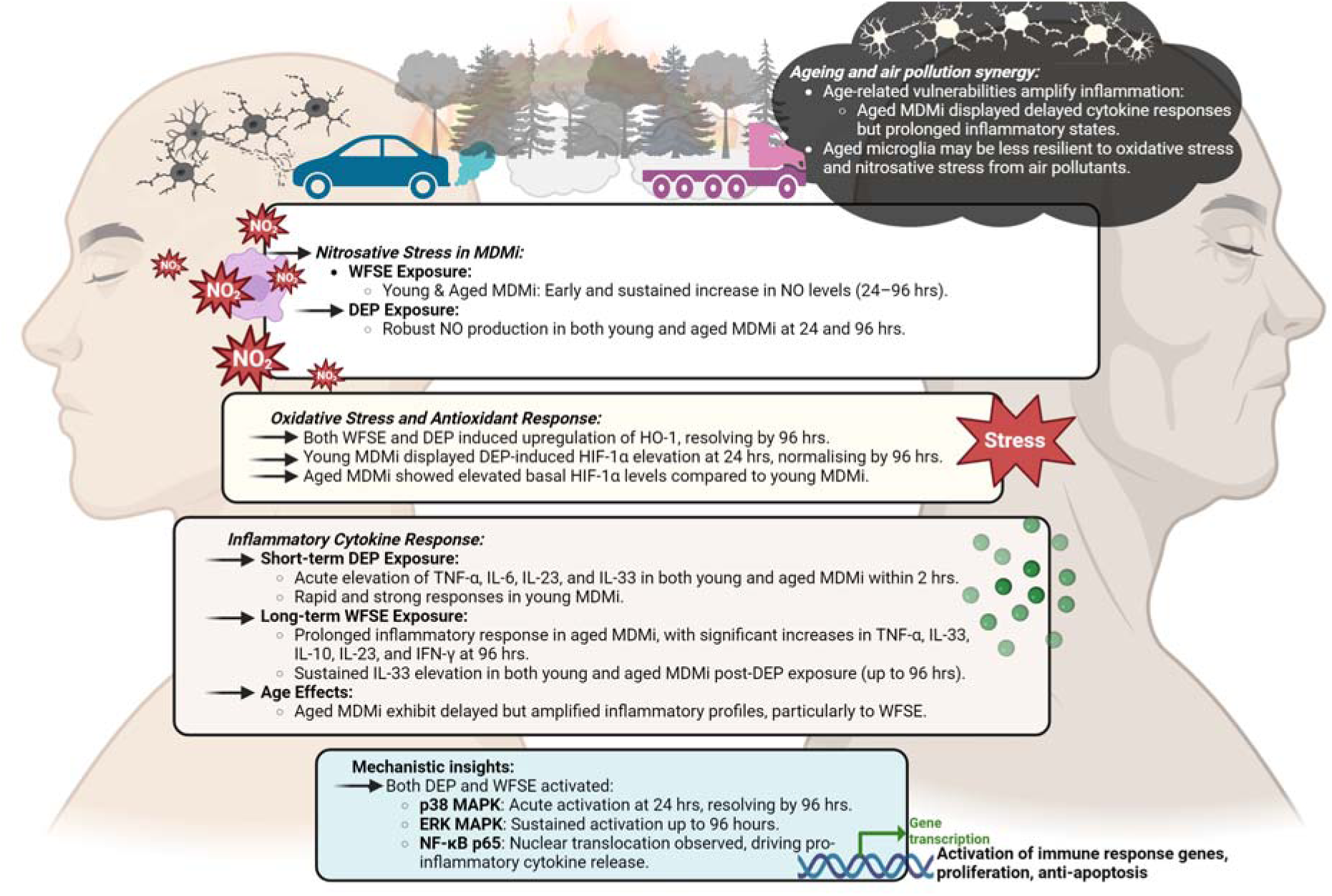
Graphical summary of key findings between age groups exposed to WFSE and DEP. WFSE and DEP exposure activate microglia, triggering inflammatory responses and nitrosative stress in both young and aged MDMi. DEP causes a stronger acute response, while WFSE induces a stronger chronic response, particularly in aged microglia. Both NF-κB and MAPK pathways are involved in this response, highlighting potential targets for mitigating air pollution’s impact on brain health, especially in the elderly.

## Acknowledgements

We thank the volunteers from QIMRB-MRI who have donated blood to this project and the elderly participants recruited through the PISA study, which is coordinated by Professor Michael Breakspear, Kerrie McAloney, Jessica Adsett and Natalie Garden. We thank the QIMR Microscopy, Sample Processing for their assistance and Dr Gunter Hartel for his advice on statistical analysis. We thank Dr Sadegh Niazi-Esfyani and Dr Priyanka Arora for providing the DEP, and Dr Jun Yan and A/Professor Judith Greer for providing NF-_K_B antibodies.

## Article Notes

### Ethics approval and consent to participate

All research adhered to the ethical guidelines on human research outlined by the National Health and Medical Research Council of Australia (NHMRC). Ethical approval was obtained from QIMR Berghofer Medical Research Institute. All participants provided informed consent before participating in the study.

### Availability of data and materials

The datasets used during the current study are available from the corresponding author upon reasonable request.

## Authors’ contributions

Conceived the study design, C.C-L, A.R.W, H.Q.

Performed experiments, analysis and interpretation of data C.C-L, M.F.N, R.S., L.A.M, F.E, Y.F.S, T.H.Y, E.V, P.F.A.M.K.L, T.L.R, Z.R, S.H, P.N.R, A.R.W, H.Q

Drafting manuscript C.C-L, R.S, A.R.W, H.Q.

Critical review and revision of manuscript C.C-L, M.F.N, R.S, T.L.R, Z.R, S.H, A.R.W, H.Q. All authors contributed to study design and approved the final manuscript.

---

**Supplementary Table 1.**

|  |  | Young Healthy donors | Aged Healthy donors |
| --- | --- | --- | --- |
| N° of donors |  | *n* = 6 | *n* = 6 |
| Sex of donors | Females (%) | 50% (3/6) | 50% (3/6) |
|  | Males (%) | 50% (3/6) | 50% (3/6) |
| Age of donors (mean ± SD) |  | 29.8 ± 3.9 | 67.4 ± 3.4 |

**Supplementary Table 2.**
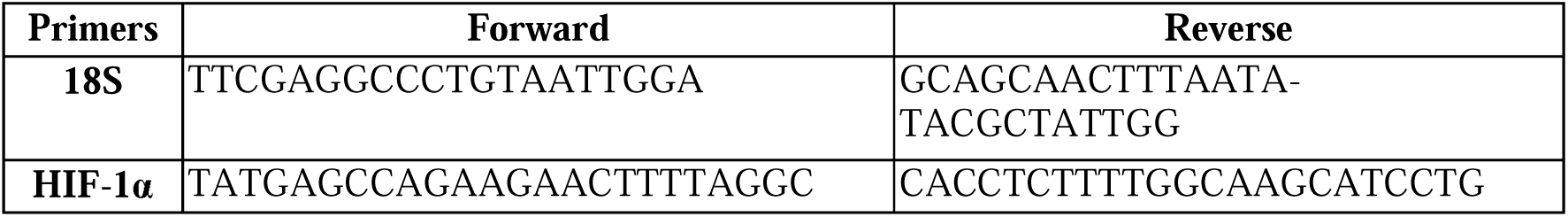

