## Supplementary Figures for "Impact of wildfire smoke and diesel exhaust on inflammatory response in aging human microglia"

^+^Co-last authors

*Corresponding:

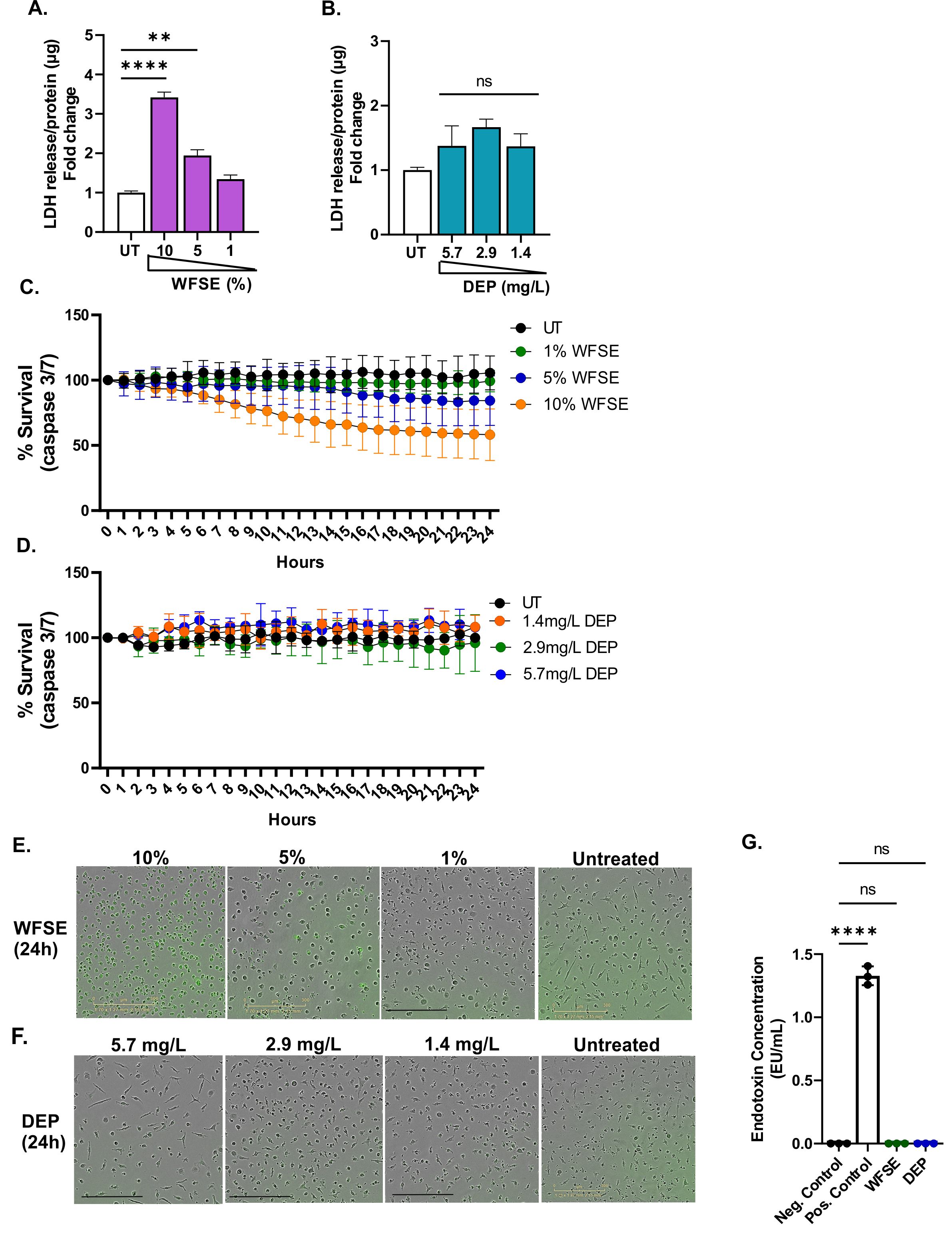

**Figure S1. Evaluation of cytotoxic and immune responses of WFSE and DEP on MDMi cells. (A)** Significant increases in LDH levels were observed in MDMi exposed to 10% and 5% WFSE after 24 hr, indicating cytotoxicity, while no significant changes were seen in **(B)** DEP-exposed MDMi at concentrations of 5.7 mg/L, 2.9 mg/L, and 1.4 mg/L. LDH was normalised against total protein for each condition and fold change to control. **(C)** Apoptosis, measured by caspase 3/7 activity, showed elevated cell death in MDMi treated with 10% (after 7 hr) and 5% WFSE (after 16 hr). **(D)** No changes in caspase 3/7 activity were observed in DEP-exposed cells. **(E)** Representative image showing caspase 3/7-positive cells in MDMi exposed to high concentrations of WFSE, **(F)** with no caspase 3/7 positivity observed in DEP-exposed MDMi. **(G)** Endotoxin testing confirmed the absence of endotoxins in both WFSE and DEP extracts. Data presented as mean ± SEM. Statistical analysis between two groups was performed using Student’s *t* test and between multiple groups using one-way ANOVA. Statistical significance indicated as follows: ****p < 0.01, ****p < 0.0001*.* Scale bars represent 300 µm.

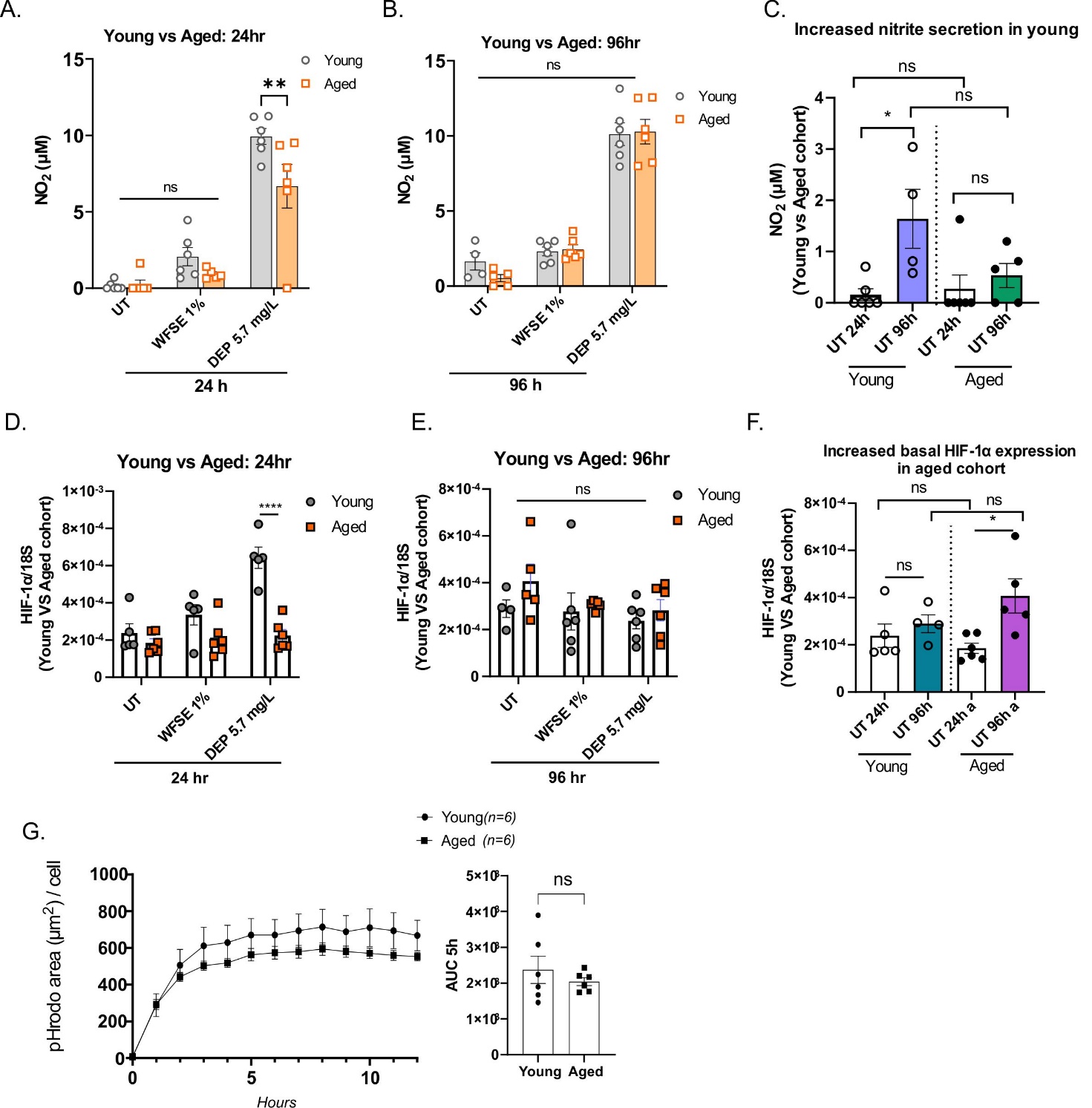

**Fig S2. WFSE and DEP induce nitrosative stress but did not alter phagocytic capability in young and aged MDMi. (A)** Increased nitrosative stress levels in young compared to aged MDMi exposed to 5.7 mg/L DEP but not 1% WFSE at 24 hr. No differences in nitrosative stress between young and aged MDMi at **(B)** 96 hr. **(C)** Basal NO levels increased in young MDMi from 24 hr to 96 hr, while aged MDMi showed no comparable increase over the same period. No significant differences observed between young and aged MDMi at each individual time point (24 hr vs. 24 hr for each cohort). **(D)** Increased HIF-1α levels in young compared to aged MDMi exposed to 5.7 mg/L DEP at 24 hr. No differences in HIF-1α levels between young and aged MDMi at **(E)** 96 hr. **(F)** Increased basal HIF-1α levels in aged MDMi at 96 hr compared to 24 hr. No significant differences observed between young and aged MDMi at each individual time point (24 hr vs. 24 hr for each cohort). **(G)** A decreasing trend in phagocytic activity was observed in aged MDMi compared to young MDMi, n = 6. Data presented as mean ± SEM. AUC = Area under curve. Statistical significance indicated as follows: ***p < 0.05, ****p < 0.0001.

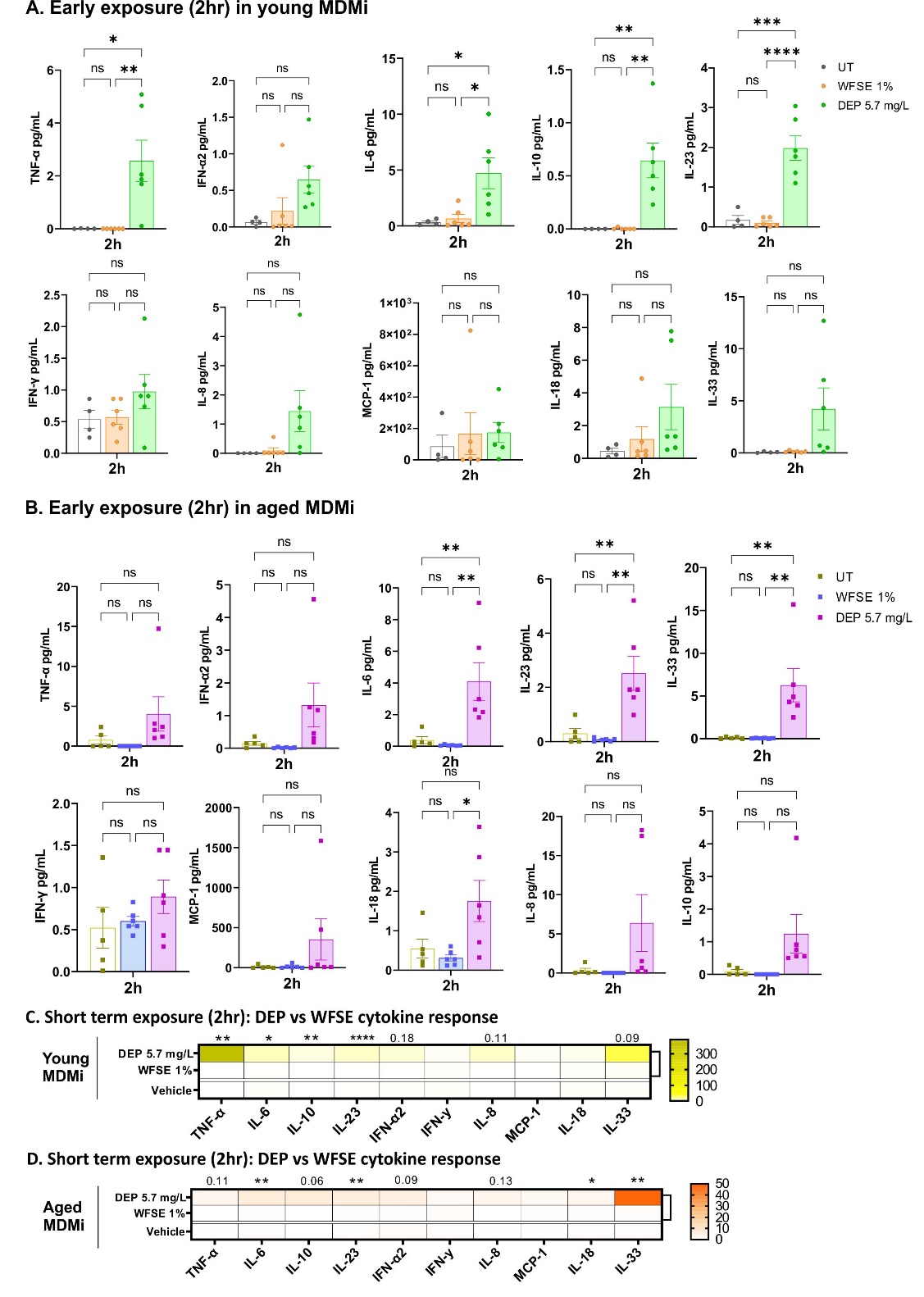

**Figure S3. Pronounced and rapid cytokine response to short-term DEP exposure compared to WFSE in both MDMi cohorts. (A)** Cytokine profiling of young MDMi exposed to 1% WFSE (orange) or 5.7 mg/L DEP (green). **(B)** Cytokine profiling of aged MDMi exposed to 1% WFSE (blue) or 5.7 mg/L DEP (magenta). **(C)** Comparison between DEP and WFSE exposure in young MDMi under short-term conditions (2 hr), shows significantly higher levels of IL-6, IL-23, IL-18, and IL-33 in cells exposed to DEP compared to WFSE. **(D)** Comparison between DEP and WFSE exposure in aged MDMi under short-term conditions, shows significantly higher levels of IL-6, IL-23, IL-18, and IL-33 in cells exposed to DEP compared to WFSE. Data presented as mean ± SEM, with data points representing biological replicates (donors). Statistical significance was determined using Student’s *t*-test for two-group comparisons and one-way or two-way ANOVA for multiple groups. Significance levels indicated as follows: *p < 0.05, **p < 0.01, ***p < 0.001, ****p < 0.0001.

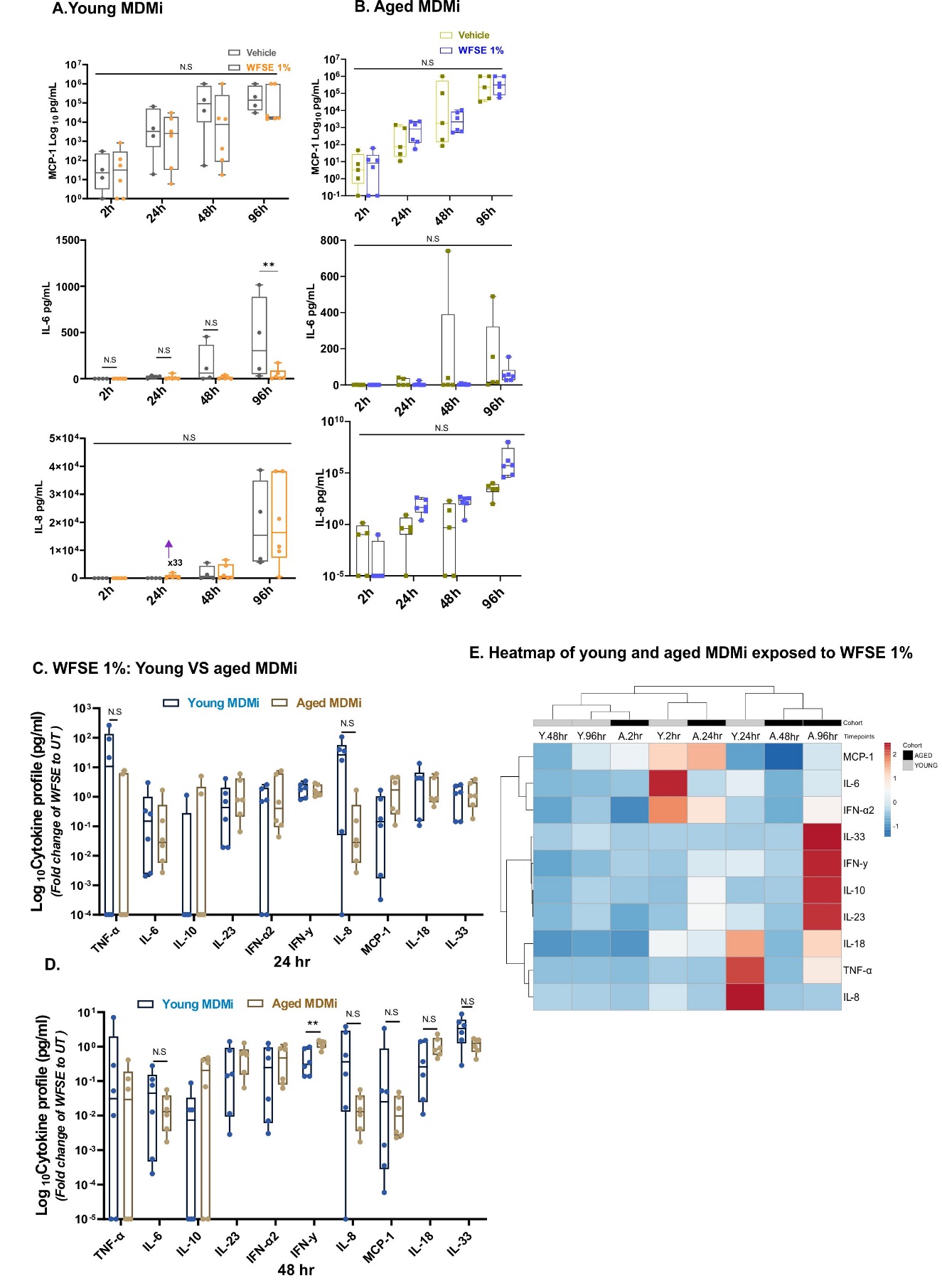

**Figure S4. Long-term exposure to WFSE 1% reveals enhanced inflammatory response in aged MDMi. (A)** Longitudinal time-course analysis of young MDMi or **(B)** aged MDMi treated with WFSE 1% or untreated control shows time-dependent kinetics of MCP-1, IL-6, and IL-8. **(C)** Comparative analysis between young and aged MDMi exposed to WFSE 1% over 24 hr or **(D)** 48hr. Cytokine changes presented as fold-change relative to the untreated control for the corresponding time point. **(E)** Heatmap of cytokine changes across young and aged MDMi exposed to WFSE 1% over 2 hr, 24 hr, 48 hr, and 96 hr. Cytokine values were log-transformed. Both rows and columns were clustered using correlation distance and average linkage**.** Data presented as mean ± SD, with data points representing biological replicates (donors; young, n = 4-6; aged, n = 5-6). Statistical significance was determined using Student’s *t*-test for two-group comparisons and one-way or two-way ANOVA for multiple groups. Significance levels indicated as follows: ****p < 0.01*.*

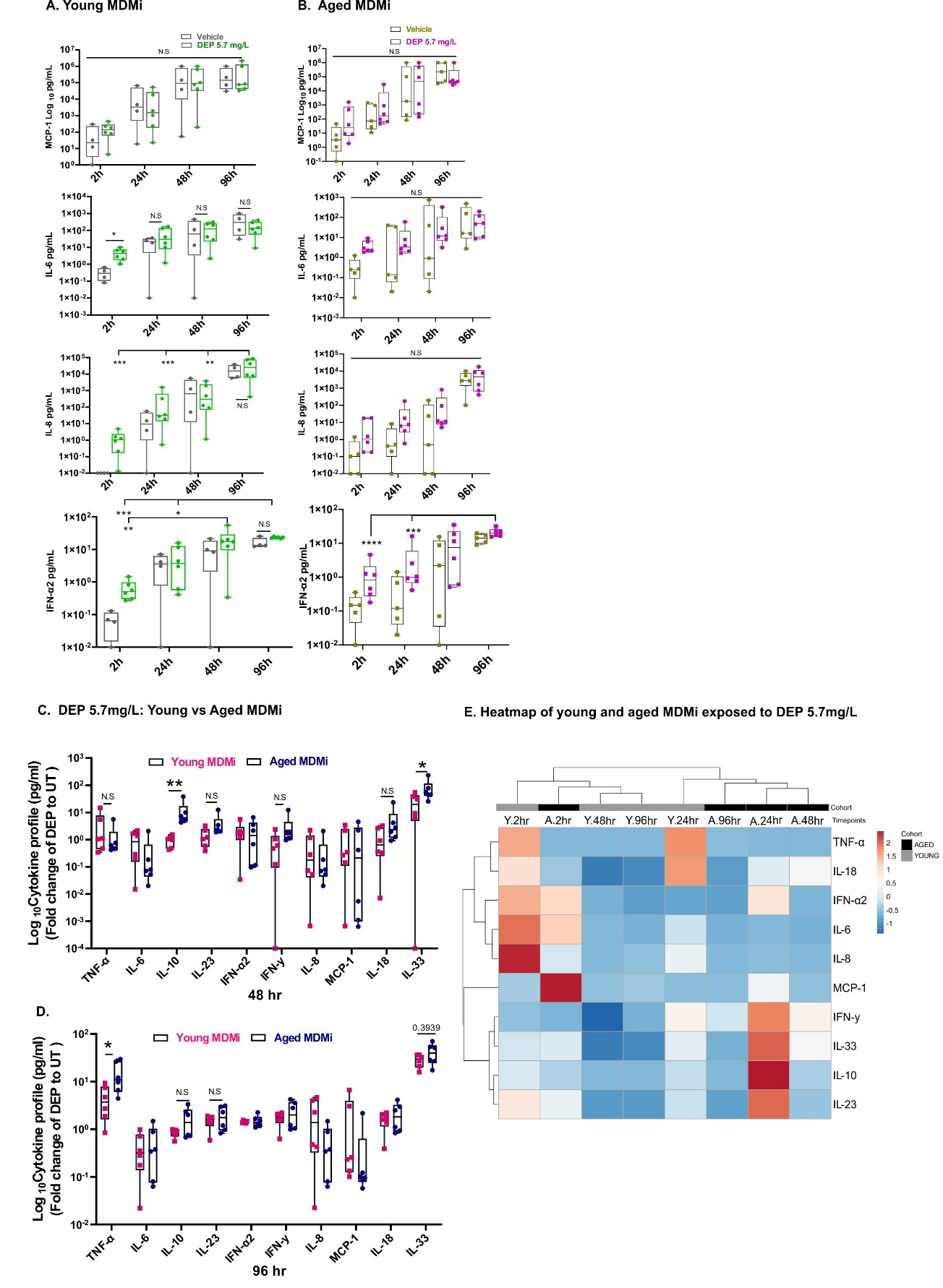

**Figure S5. Aged MDMi exhibit peak cytokine response at 24 hr following DEP 5.7 mg/L** **exposure. (A)** Longitudinal time-course analysis of young MDMi or **(B)** aged MDMi treated with DEP 5.7 mg/L or untreated control shows time-dependent kinetics of MCP-1, IL-6, IL-8, IFN-α2. **(C)** Comparative analysis between young and aged MDMi exposed to WFSE 1% over 48 hr or **(D)** 96hr. Cytokine changes presented as fold-change relative to the untreated control for the corresponding time point. **(D)** Heatmap of cytokine changes across young and aged MDMi exposed to DEP 5.7 mg/L over 2 hr, 24 hr, 48 hr, and 96 hr. Cytokine values were log-transformed. Both rows and columns were clustered using correlation distance and average linkage**.** Data presented as mean ± SD, with data points representing biological replicates (donors; young, n = 4-6; aged, n = 5-6). Statistical significance determined using Student’s *t*-test for two-group comparisons and one-way or two-way ANOVA for multiple groups. Significance levels indicated as follows: *p < 0.05, **p < 0.01, ***p < 0.001, ****p < 0.0001.

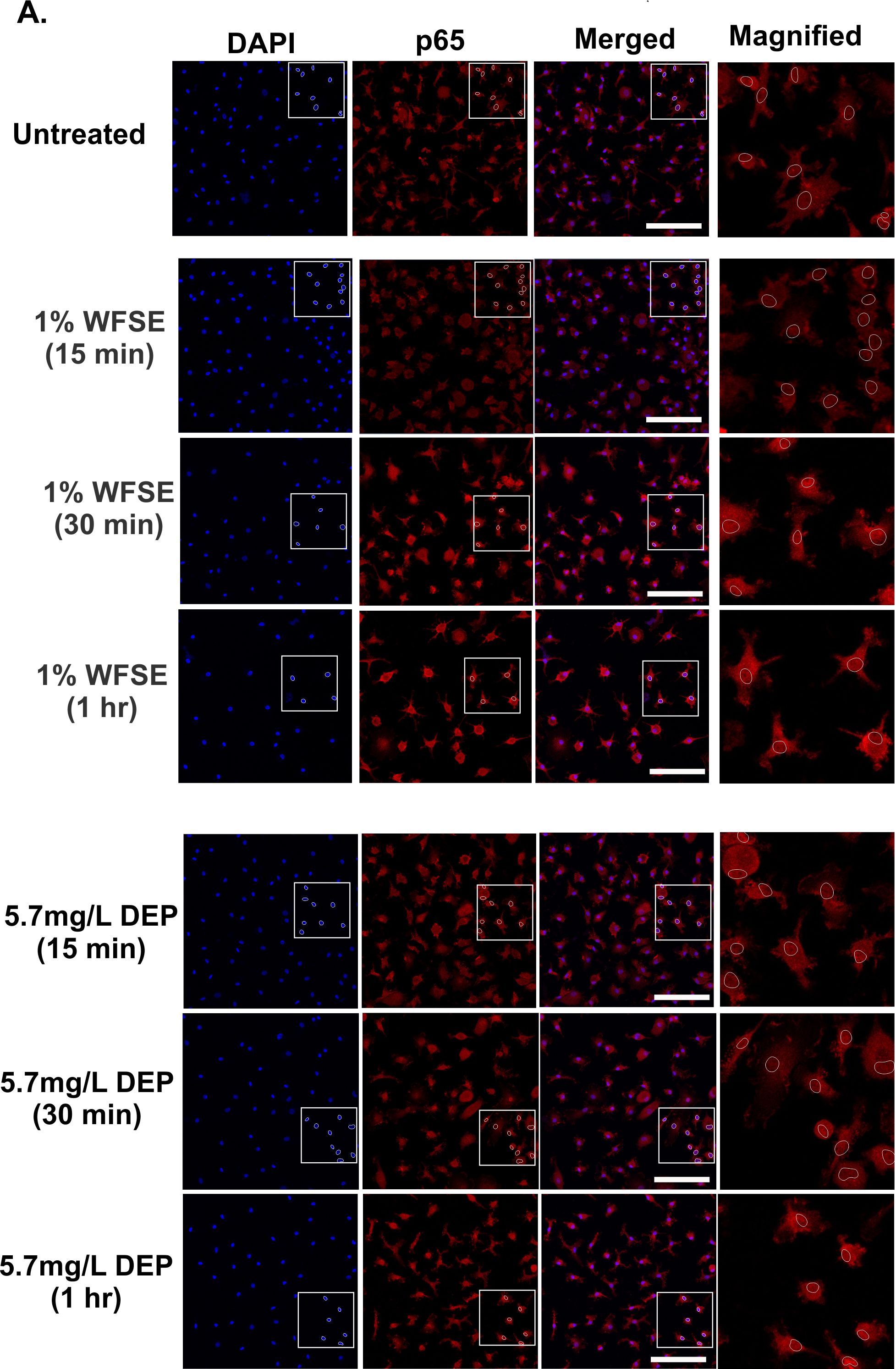

**Figure S6.** **Lack of p65 activation in MDMi exposed to air pollutants at earlier time points.** MDMi were exposed to either 1% WFSE or 5.7 mg/L DEP for 15 min, 30 min, and 1 hr. No nuclear translocation of p65 was observed at these earlier time points; total n = 50 cells analysed from n = 3 donors. Magnified images show the areas within the white boxes. Dotted circles in the panels indicate nuclear localisation as marked by DAPI staining. Scale bars represent 50 µm.
